# Hidden GPCR structural transitions addressed by multiple walker supervised molecular dynamics (mwSuMD)

**DOI:** 10.1101/2022.10.26.513870

**Authors:** Giuseppe Deganutti, Ludovico Pipitò, Roxana M. Rujan, Tal Weizmann, Peter Griffin, Antonella Ciancetta, Stefano Moro, Christopher A. Reynolds

**Affiliations:** Centre for Health and Life Sciences, Coventry University, Alison Gingell Building, Coventry, CV1 5FB, UK; Dipartimento di Scienze Chimiche, Farmaceutiche ed Agrarie, University of Ferrara, Via Fossato di Mortara 17/19, 44121 Ferrara, Italy; Molecular Modeling Section (MMS), Dipartimento di Scienze del Farmaco, University of Padua via Marzolo 5, 35131, Padova, Italy; School of Life Sciences, University of Essex, Wivenhoe Park, Colchester, CO4 3SQ

**Keywords:** G protein-coupled receptors, G protein, binding, protein activation, molecular dynamics, supervised molecular dynamics, GDP release

## Abstract

The structural basis for the pharmacology of G protein-coupled receptors (GPCRs), the most abundant membrane proteins and the target of about 35% of approved drugs, is still a matter of intense study. What makes GPCRs challenging to study is the inherent flexibility and the metastable nature of interaction with extra-and intracellular partners that drive their effects. Here, we present a molecular dynamics (MD) adaptive sampling algorithm, namely multiple walker supervised molecular dynamics (mwSuMD), to address complex structural transitions involving GPCRs without energy input. We first report the binding and unbinding of the vasopressin peptide from its receptor V_2_. Successively, we present the complete transition of the glucagon-like peptide-1 receptor (GLP-1R) from inactive to active, agonist and G_s_-bound state, and the GDP release from G_s_. To our knowledge, this is the first time the whole sequence of events leading from an inactive GPCR to the GDP release is simulated without any energy bias. We demonstrate that mwSuMD can address complex binding processes intrinsically linked to protein dynamics out of reach of classic MD.

## Introduction

Supervised molecular dynamics^1,2^ (SuMD) is an efficient adaptive sampling technique for studying ligand-receptor binding and unbinding pathways; here, we present the multiple walker enhancement (mwSuMD) to study a significantly wider range of structural transitions relevant to the drug design. We validated the method by applying it to G protein-coupled receptors (GPCRs), since their inherent flexibility is essential to their function and because these are the most abundant family of membrane receptors in eukaryotes^3^ and the target for more than one-third of drugs approved for human use^4^.

Vertebrate GPCRs are subdivided into five subfamilies (Rhodopsin or class A, Secretin or class B, Glutamate or class C, Adhesion, and Frizzled/Taste2) according to function and sequence^5,6^. X-ray and cryo-electron microscopy (cryo-EM) show that GPCRs possess seven transmembrane (TM) helices connected by three extracellular loops (ECLs) and three intracellular loops (ICLs), with an extended and structured N-terminal extracellular domain (ECD) in all subtypes, but class A. The primary function of GPCRs is transducing extracellular chemical signals into the cytosol by binding and activating four G protein families (G_s/olf_, G_i/o_, G_12/13_, and G_q/11_) responsible for decreasing (G_i/o_) or increasing (G_s/olf_) cyclic adenosine-3’,5’- monophosphate (cAMP), and generating inositol-1,4,5-triphosphate (IP_3_) and diacylglycerol (DAG) to increase Ca^2+^ intracellular levels (G_q_)^7^.

Many aspects of GPCR pharmacology remain elusive: for example, the structural determinants of the selectivity displayed towards specific G proteins or the ability of certain agonists to drive a preferred intracellular signaling pathway over others (*i.e.* functional selectivity or bias)^8^. GPCRs are challenging proteins to characterize experimentally due to their inherent flexibility and the transitory nature of the complexes formed with extracellular and intracellular effectors. Importantly, agonists can allosterically modify the receptor selectivity profile by imprinting unique intracellular conformations from the orthosteric binding site. The mechanism behind these phenomena is one of the outstanding questions in the GPCR field^9^.

Molecular dynamics (MD) is a powerful computational methodology that predicts the movement and interactions of (bio)molecules in systems of variable complexity, at atomic detail. However, classic MD sampling is limited to the microsecond or, in the best conditions, the millisecond time scale^10,11^. For this reason, different algorithms have been designed to speed up the simulation of rare events such as ligand (un)binding and conformational transitions. Amongst the most popular and effective ones are metadynamics^12^ and path collective variables metadynamics^13^, accelerated MD (aMD)^14^, and Gaussian-accelerated MD (GaMD)^15^, which introduce an energy potential to overcome the energy barriers preventing the complete exploration of the free energy surface, thus *de facto* bias the simulation. Energetically unbiased MD protocols, on the other hand, comprise weighted ensemble MD (weMD)^16^, swarms approach^17^, AdaptiveGoal^18^, and SuMD^1,19^, which have largely been applied to (unbinding) small molecules, peptides, and small proteins^1,19–23^. Since SuMD is optimized for (un)bindings, we have designed mwSuMD to address more complex conformational transitions and protein-protein associations. GPCRs preferentially couple to very few G proteins out of 23 possible counterparts^9,24^. It is increasingly accepted that dynamic and transient interactions determine whether the encounter between a GPCR and a G protein results in productive or unproductive coupling^25^. MD simulations are considered a useful orthogonal tool for providing working hypotheses and rationalizing existing data on G protein selectivity. However, so far, it has not delivered as expected. Attempts have usually employed energetically biased simulations, have been confined to the Gα subunit, or considered a pre-formed GPCR:G protein complex^24,26–28^.

Firstly, we validated mwSuMD on the nonapeptide arginine vasopressin (AVP) by simulating binding (dynamic docking) and unbinding paths from the vasopressin 2 receptor (V_2_R). Dynamic docking, although more computationally demanding than standard molecular docking, provides insights into the binding mode of ligands in a fully hydrated and flexible environment. Moreover, it informs about binding paths and the complete mechanism of formation leading to an intermolecular complex, delivering in the context of binding kinetics^29^ and structure-kinetics relationship (SKR) studies^30^. We then studied the class B1 GPCR glucagon-like peptide-1 receptor (GLP-1R) activation by the small molecule PF06882961. GLP-1R is a validated target in type 2 diabetes and probably the best-characterized class B1 GPCR from a structural perspective. GLP-1R is the only class B1 receptor with structurally characterized non-peptidic orthosteric agonists, which makes it a model system for studying the druggability of the entire B1 subfamily. After GLP-1R agonist binding and activation, the coupling of G_s_ and the release of GDP, the rate-limiting step of the G protein activation, was simulated for the first time using an energy-unbiased method.

These results demonstrate the usefulness of mwSuMD for illuminating the molecular events involved in GPCR function.

## Results and Discussion

### Short mwSuMD time windows improve the AVP dynamic docking prediction

Arginine vasopressin (AVP) is an endogenous hormone (**Figure S1a)** that mediates antidiuretic effects on the kidney by signaling through three class A GPCR subtypes: V_1a_ and V_1b_ receptors activate phospholipases via G_q/11_, while the V_2_ receptor (V_2_R) activates adenylyl cyclase by interacting with G_s_ ^31^ and is a therapeutic target for hyponatremia, hypertension, and incontinence^32^. AVP is amphipathic and in the bound state interacts with both polar and hydrophobic V_2_R residues located on TM helices and ECLs (**Figure S1b**). Although AVP presents an intramolecular C1-C6 disulfide bond that limits the backbone’s overall conformational flexibility, it has many rotatable bonds, making dynamic docking complicated^33^. We compared the performance of mwSuMD to the parent algorithm SuMD in reconstructing the experimental V_2_R:AVP complex using different settings, simulating a total of 92 binding events (**Table S1**). As a reference, the AVP RMSD during a classic (unsupervised) equilibrium MD simulation of the X-ray AVP:V_2_R complex was 3.80 ± 0.52 Å (**Figure S2**). SuMD^1,19^ produced a minimum root mean square deviation (RMSD) to the cryo-EM complex of 4.28 Å, with most of the replicas (*i.e.,* distribution’s mode) having an RMSD close to 10 Å (**Figure S3a**). mwSuMD, with the same settings (**Figure S3b, Table S1**) in terms of time window duration (600 ps), metric supervised (the distance between AVP and V_2_R), and acceptance method (slope) produced slightly more precise results (i.e., distribution’s mode RMSD = 7.90 Å) but similar accuracy (minimum RMSD = 4.60). Supervising the AVP RMSD to the experimental complex rather than the distance (**Figure S3c**) and using the SMscore (**Equation 1**) as the acceptance method (**Figure S3d**) worsened the prediction. Supervising distance and RMSD at the same time (**Figure S3e**), employing the DMscore (**Equation 2**), recovered accuracy (minimum RMSD = 4.60 Å) but not precision (distribution mode RMSD = 12.40 Å). Interestingly, decreasing the time window duration from 600 ps to 100 ps impaired the SuMD ability to predict the experimental complex (**Figure 1a**), but enhanced mwSuMD accuracy and precision (**Figure 1b-d**). The combination of RMSD as the supervised metric and SMscore produced the best results in terms of minimum RMSD and distribution mode RMSD, 3.85 Å and 4.40 Å, respectively (**Figure 1d, Video S1**), in agreement with the AVP deviations in the equilibrium MD simulation of the X-ray AVP:V_2_R complex (**Figure S2**). These results confirm the inherent complexity of reproducing the AVP:V_2_R complex via dynamic docking and suggest that short time windows can improve mwSuMD performance on this system. However, it is necessary to know the final bound state to employ the RMSD as the supervised metric, while the distance is required to dynamically dock ligands with unknown bound conformation as previously reported^1,24^. Both distance and RMSD-based simulations delivered insights into the binding path and the residues involved along the recognition route. For example, mwSuMD pinpointed V_2_R residues E184^ECL2^, P298^ECL3^, and E303^ECL3^ (**Figure S4a**) as involved during AVP binding, although not in contact with the ligand in the orthosteric complex. None of them are yet characterized through mutagenesis studies according to the GPCRdb^34^.

**Figure 1.**
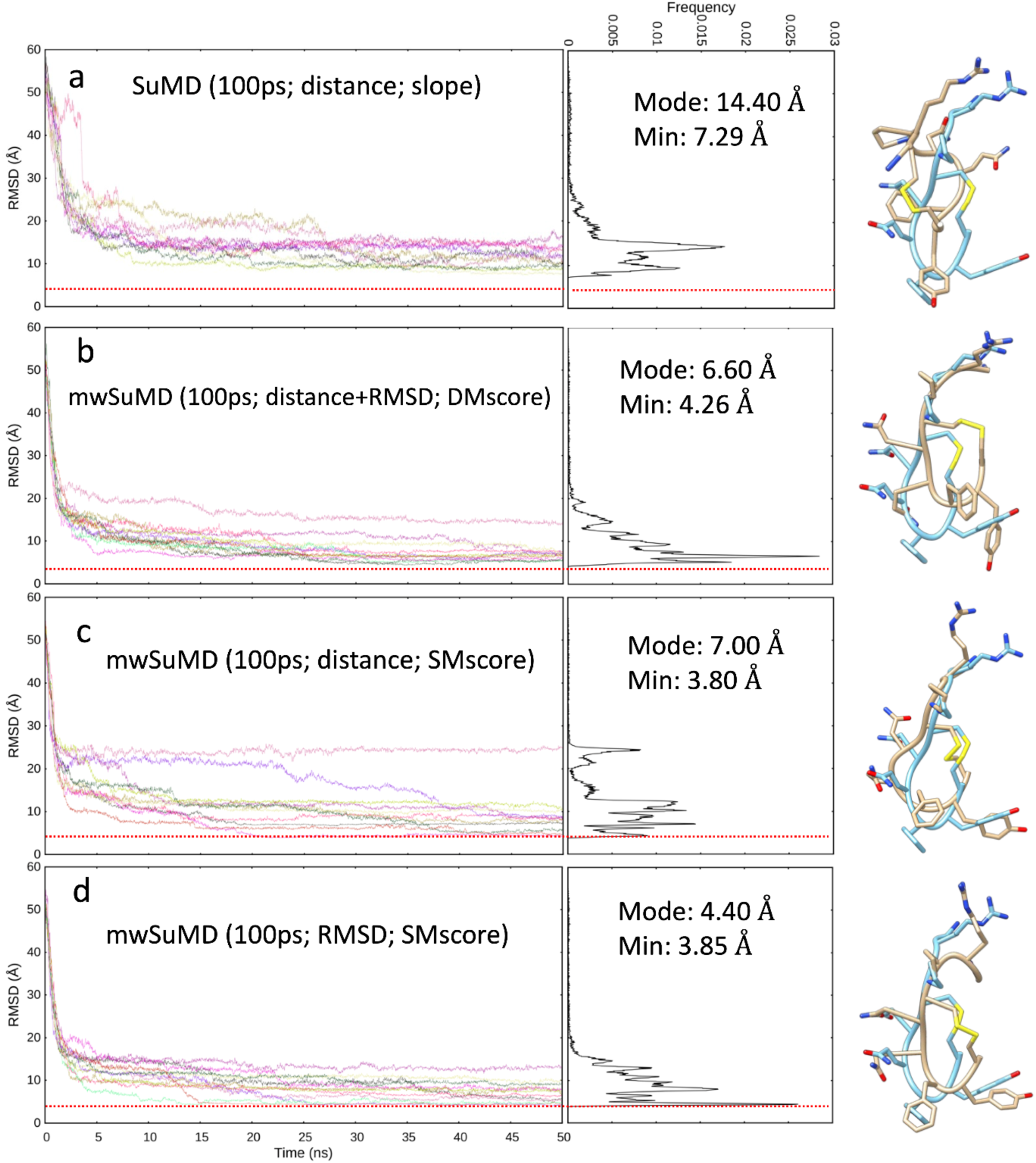
AVP SuMD and mwSuMD binding simulations to V_2_R (100 ps time windows). For each set of settings **(a-d)** the RMSD of AVP Cα atoms to the cryo-EM structure 7DW9 is reported during the time course of each SuMD (**a**) or mwSuMD (**b-d**) replica alongside the RMSD values distribution and the snapshot corresponding to the lowest RMSD values (AVP from the cryo-EM structure 7DW9 is in a cyan stick representation, while AVP from simulations is in a tan stick representation). A complete description of the simulation settings is reported in Table S1 and the Methods section. The dashed red line indicates the AVP RMSD during a classic (unsupervised) equilibrium MD simulation of the X-ray AVP:V_2_R complex (**Figure S2**).

Further to binding, a SuMD approach was previously employed to reconstruct the unbinding path of ligands from several GPCRs^2,35^. We assessed mwSuMD’s capability to simulate AVP unbinding from V_2_R. Five mwSuMD and five SuMD replicas were collected using 100 ps time windows (**Table S1**). Overall, mwSuMD outperformed SuMD in terms of time required to complete a dissociation (**Figure S5, Video S2**), producing dissociation paths almost 10 times faster than SuMD. Such rapidity in dissociating inherently produces a limited sampling of metastable states along the pathway, which can be compensated by seeding classic (unsupervised) MD simulations from configurations extracted from the unbinding pathway^36,37^. Here, the V_2_R residues involved during the dissociation were comparable to the binding (**Figure S4b**), although ECL2 and ECL3 were slightly more involved during the association than the dissociation, in analogy with other class A and B GPCRs^21,36^.

### PF06882961 binding and GLP-1R activation

The GLP-1R has been captured by cryo-electron microscopy (cryo-EM) in both the inactive and the active (G_s_-bound) conformations and in complex with either peptide or non-peptide agonists^38–43^. In the inactive GLP-1R, residues forming the binding site for the non-peptide agonist PF06882961 are dislocated and scattered due to the structural reorganization of the transmembrane domain (TMD) and extracellular domain (ECD) (**Figure S6**) that occurs on activation. Moreover, GLP-1R in complex with GLP-1 or different agonists present distinct structural features, even amongst structurally related ligands (**Figure S7**). This complicates the scenario and suggests divergent recognition mechanisms amongst different agonists. We simulated the binding of PF06882961, reaching an RMSD to its bound conformation in 7LCJ of 3.79 ± 0.83 Å (computed on the second half of the merged trajectory, superimposing on GLP-1R Cα atoms of TMD residues 150 to 390), using multistep supervision on different system metrics (**Figure 2**) to model the structural hallmark of GLP-1R activation (**Video S5**, **Video S6**).

**Figure 2.**
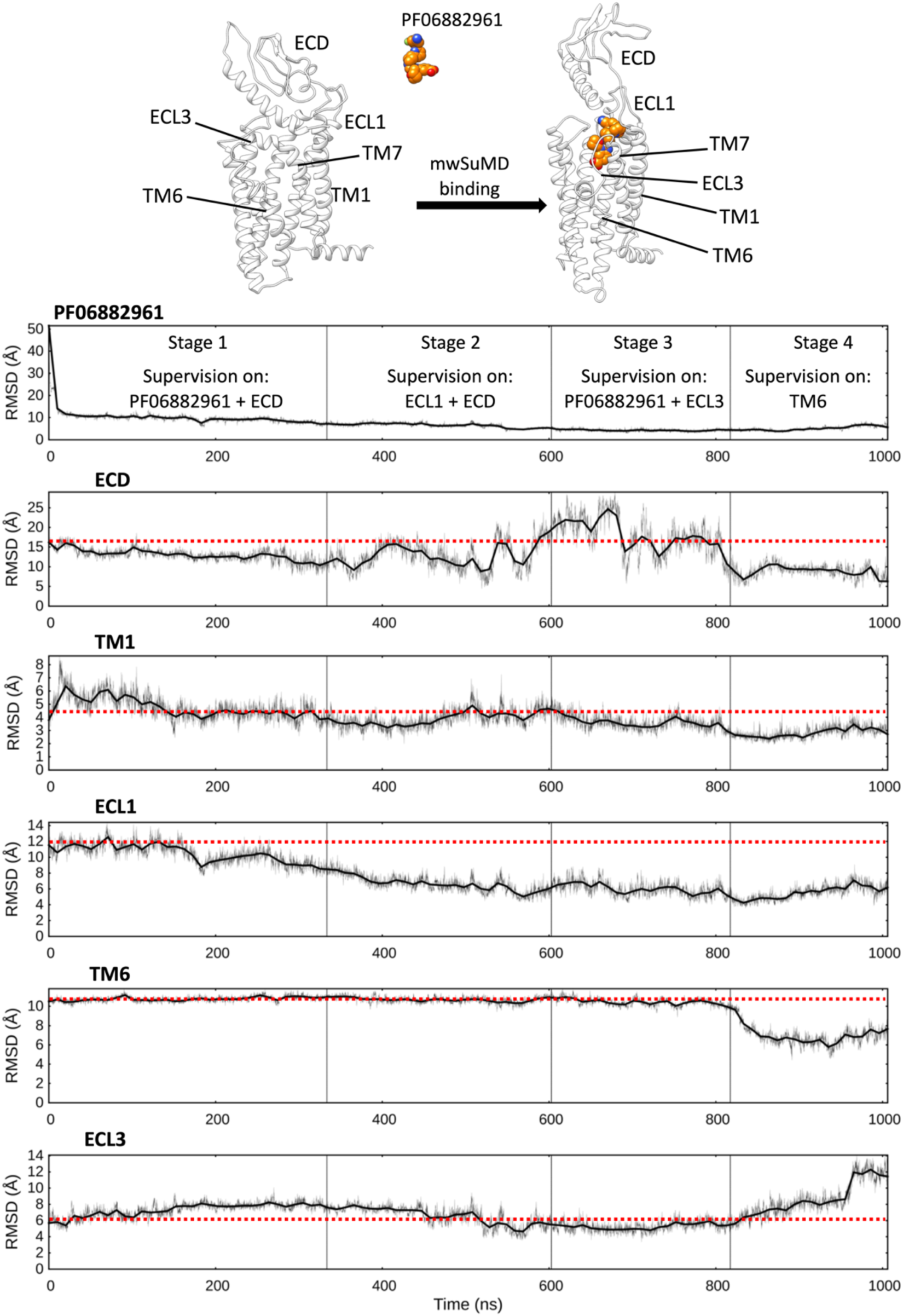
mwSuMD simulation of PF06882961 binding to GLP-1R and receptor activation. Each panel reports the root mean square deviation (RMSD) to the position of the ligand in the active state (top panel) or a GLP-1R structural element over the time course (all but ECL3 converging to the active state). ECD: extracellular domain; TM: transmembrane helix; ECL: extracellular loop. The mwSuMD simulation was performed with four different settings over 1 microsecond in total. The red dashed lines show the initial RMSD value for reference.

Several metrics were supervised consecutively. Firstly, the distance between PF06882961 and the TMD as well as the RMSD of the ECD to the active state (stage 1); secondly, the RMSD of ECD and ECL1 to the active state (stage 2); thirdly, the RMSD of PF06882961 and ECL3 to the active state (stage 3); lastly, only the RMSD of TM6 (residues I345-F367, Cα atoms) to the active state (stage 4). The combination of these supervisions produced a conformational transition of GLP-1R towards the active state (**Figure 2, Video S6**). Noteworthy, mwSuMD, like any other CV-based technique, requires some knowledge of the simulated system. The sequence of these supervisions was arbitrary and does not necessarily reflect the right order of the steps involved in GLP-1R activation. This kind of planned multistep approach is feasible when the end-point receptor inactive and active structures are available, and the inherent flexibility of different domains is known. In class B GPCRs, the ECD is the most dynamic sub-structure, followed by the ECL1 and ECL3, which display high plasticity during ligand binding^21,44^. For this reason, we first supervised these elements of GLP-1R, leaving the bottleneck of activation, TM6 outward movement, as the last step. However, the protocol employed can be tweaked to study how each conformational transition takes place and influences the receptor domains. Structural elements not directly supervised, such as TM1 or TM7, were influenced by the movement of supervised helixes or loops and therefore displayed an RMSD reduction to the active state. For example, the supervision of ECL3 (stage 3) and TM6 (stage 4) facilitated the spontaneous rearrangement of the ECD to an active-like conformation after the ECD had previously experienced transient high flexibility during stages 2 and 3 (**Figure 2**). These results suggest a concerted conformational transition for ECD and ECL1 during the binding of PF06882961 and an allosteric effect between ECL3 and the bottom of TM6. While the intracellular polar interactions were destabilized by the ECL3 transition to an active-like conformation (stages 2 and 3), the outward movement of TM6 (stage 4) did not favor the closure of ECL3 towards PF06882961, which appears to be driven by direct interactions between the ligand and R310^5.40^ or R380^7.35^. Interestingly, the mwSuMD simulation during Stage 4 (TM6 supervision) sampled a counterclockwise helix rotation (**Figure S8a**) consistent with the GLP-1R cryo-EM structures in the active, G_s_-coupled state^41,45^.

It is worth noting that 6LN2, the only inactive GLP-1R structure available complete with the ECD, was stabilized and solved with an antibody bound to the ECD. This strategy might have forced the ECD into a closed conformation that engages the EC *vestibule* of GLP-1R and possibly restrained the whole TMD in an altered conformation that deviates from the physiological conditions. This might explain why the RMSD of the TMD elements monitored during the simulation rarely reach values lower than 3 or 4 Å.

During the supervision of ECL3 and PF06882961 (stage 3), we observed a loosening of the intracellular polar interactions that stabilize GLP-1R TM6 in the inactive state. As a result, the subsequent supervision of TM6 (residues I345-F367, Cα atoms) rapidly produced the outward movement of TM6 towards the active state, in the last step of the mwSuMD simulation (stage 4). A more detailed analysis revealed that the central polar network, which is pivotal for mediating GLP-1 signalling^46^, and the residues at the TM6 kink level adopted active-like conformations during the final stage of the simulation (**Figure 3a**). In particular, the central polar network (E364^6.53^, H363^6.52^, and Q394^7.49^) experienced side chain rearrangements (**Figure 3b**) while extensive TM6 kink dislocation occurred at L360^6.49^, P358^6.48^, L357^6.47^ and I357^6.46^ (**Figure 3c**).

**Figure 3.**
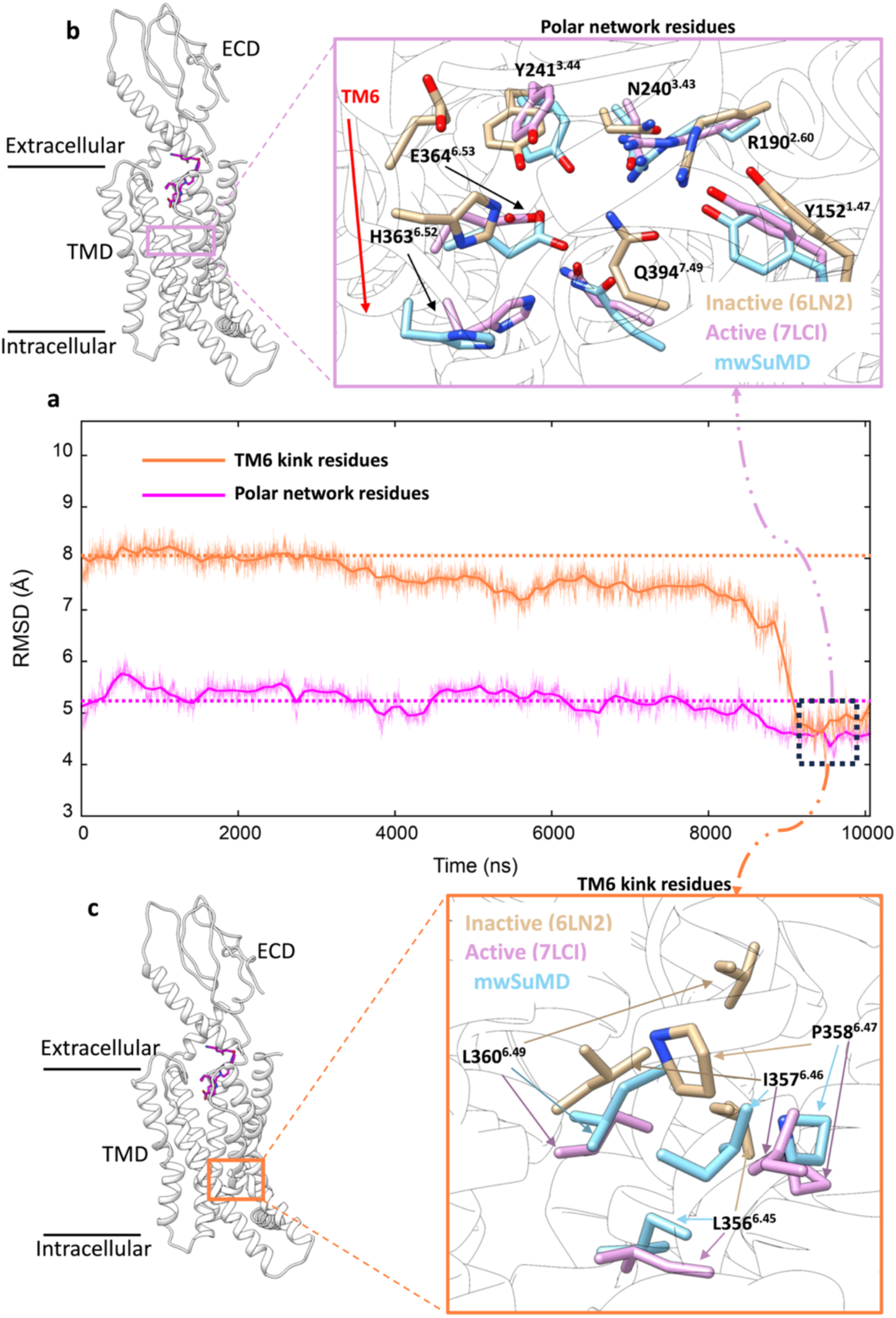
GLP-1R key structural motifs during mwSuMD GLP-1R activation. **a)** RMSD to the active state GLP-1R (7LCI) of the residues forming the central polar network (magenta) and TM6 kink (orange) during mwSuMD of receptor activation; at the end of the simulations minimum values were reached (dashed square). **b)** The position of the polar network within the core of TMD (left-hand panel) and comparison between inactive, active and mwSuMD final states for the side chains of the residues forming of the polar network; **c)** The position of the TM6 kink (right-hand panel) and comparison between inactive, active and mwSuMD final states for the side chains of the residues forming the TM6 kink.

### G_s_ protein binding to GLP-1R and GDP release

We then focused on simulating the G_s_ binding to GLP1-R, after activation, *without energy input*. For this purpose, we first tested the binding between the prototypical class A receptor β_2_ adrenoreceptor (β_2_ AR), and the stimulatory G protein (G_s_) (**Video S3, Figure S9a,b**) by supervising the distance between G_s_ helix 5 (α5) and β_2_AR as well as the RMSD of the intracellular end of TM6 to the fully active state of the receptor (see Supplementary Methods). During two out of three replicas, both Gα and Gβ achieved distance values close to 5 Å (minimum RMSD = 3.94 Å and 3.96 Å respectively), in good agreement with the reference (the β_2_ AR:G_s_ complex, PDB 3SN6, **Figure S9c).** A possible pitfall is that G proteins bear potential palmitoylation and myristoylation sites that anchor the inactive trimer to the plasma membrane^47,48^, *de facto* restraining possible binding paths to the receptor. To address this point and test different conditions, we prepared the adenosine A1 receptor (A_1_R), and its principal effector, the inhibitory G protein (G_i_) considering the G_iα_ residue C3 and G_γ_ residue C65 as palmitoylated and geranylgeranylated respectively and hence inserted in the membrane. Recently, the G_i_ binding to A_1_R was simulated by combining the biased methods GaMD with SuMD^49^ but without considering membrane-anchoring post-translational modifications. Both classic (unsupervised) and mwSuMD simulations were performed on this system for comparison (**Video S4, Figure S9d)**. In about 50 ns of mwSuMD, the G_iα_ subunit engaged its intracellular binding site on A_1_R and formed a complex in good agreement with the cryo-EM structure (PDB 6D9H, RMSD ≈ 5 Å). For comparison, 1 μs of cMD did not produce a productive engagement as the G_iα_ remained at RMSD values > 40 Å (**Figure S9d**), suggesting the effectiveness of mwSuMD in sampling G protein binding rare events without the input of energy. The membrane anchoring affected the overall G_i_ binding and the final complex, which was rotated compared to the experimental structure due to the lipidation of G_iα_ and G_γ_ (**Figure S9e)**.

Encouraged by results obtained on G_s_ and G_i_ binding to β_2_ AR and A_1_R (**Figure S9**), we extracted the GLP-1R active conformation described above and simulated the G_s_ binding to its intracellular side. Starting from the inactive, membrane-anchored G_s_, we performed three independent mwSuMD replicas by supervising the distance between G_s_ helix *α*5 and GLP-1R residues located at the intracellular binding interface (**Figure 4a,e**). All three mwSuMD replicas showed the G_s_ approaching GLP-1R, with two out of three reaching an RMSD of the G_sα_ subunit close to or less than 10 Å, compared to the experimental complex 7LCI (**Figure 4e**). Replica 2, in particular, reproduced the cryo-EM GLP-1R:G_s_ complex with RMSD values to 7LCI of 7.59 ± 1.58 Å, 12.15 ± 2.13 Å, and 13.73 ± 2.24 Å for G*_α_,* G_β_ and G_γ_, respectively. Such values do not support convergence with the static experimental structure but are not far from the RMSDs measured in our previous simulations of GLP-1R in complex with G_s_ and GLP-1^50^ (G*_α_* = 6.18 ± 2.40 Å; G_β_ = 7.22 ± 3.12 Å; G_γ_ = 9.30 ± 3.65 Å), which indicates overall higher flexibility of G_β_ and G_γ_ compared to G*_α_*, which acts as a sort of fulcrum bound to GLP-1R.

**Figure 4.**
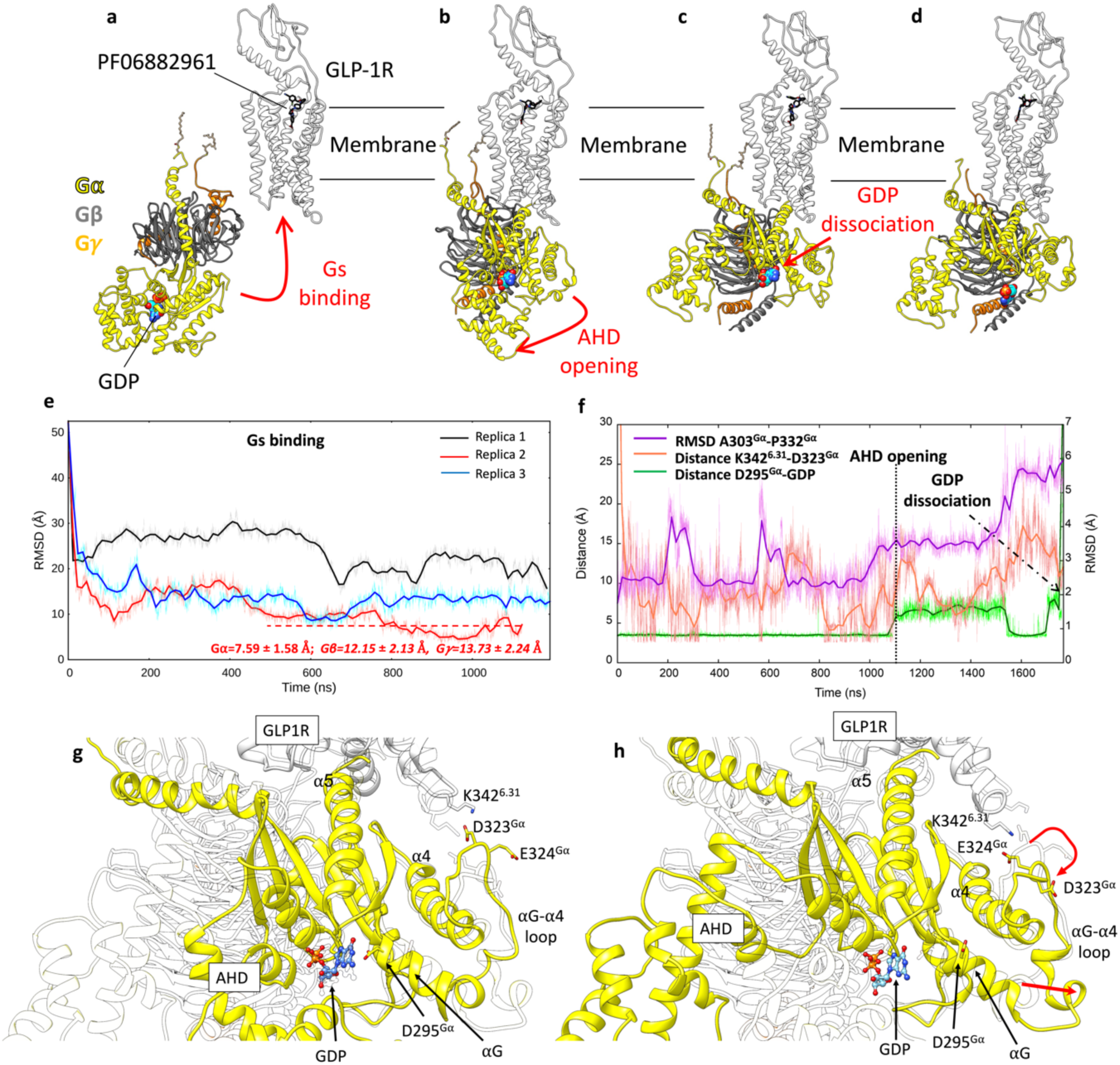
GLP-1R activation and G_s_ binding. a-d) sequence of simulated events during the mwSuMD G_s_:GLP-1R simulations. **e)** RMSD of G_s_α to the experimental GLP-1R:G_s_ complex (PDB 7LCJ) during three mwSuMD replicas; the RMSD to the experiential bound conformation (7LCJ) during the second part of Replica2 (red dashed line) is reported for each G_s_ subunit. RMSDs were computed on G_α_ residues 11 to 43 and 205 to 394 to the experimental structure 7LCI after superimposition on GLP-1R residues 140-240 Cα atoms. **f**) RMSD of the αG-α4 loop (purple), the distance between K342^6.31^ and D323^Gα^ (salmon), and the distance between GDP and D295^Gα^ (green) during G_s_ binding, AHD opening and GDP dissociation; **g)** and **h)** comparison between states extracted from before and after AHD opening. Before AHD opening (**a**), GLP-1R ICL3 interacted with D323^Gα^ and D295 ^Gα^ interacted with GDP; after AHD opening (**b**) and αG-α4 loop reorganization (curved red arrow), αG and D295 ^Gα^ moved away from GDP (straight red arrow), destabilizing its binding to G_s_.

According to the model of G protein activation, the G protein binding has the effect of allosterically stabilizing the orthosteric agonist in complex with the receptor^50^ and destabilizing the guanosine-diphosphate (GDP) bound to Gα, triggering its release and exchange with guanosine-triphosphate (GTP)^51^, upon opening of the G protein alpha-helical domain (AHD). Following this model, PF06882961 and GDP were respectively stabilized and destabilized during the simulated G_s_ association (**Figure S8b,c**). The analysis of atomic contacts along the binding path of G_s_ to GLP-1R highlights a few persistent interactions not observed in the equilibrium MD simulations of GLP-1R:G_s_ cryo-EM complexes^50^; for example, we propose the hidden interaction between D344^6.33^ and R385^α^^5^ to be important for G_s_ coupling (**Table S2**), which would explain the GLP-1 EC_50_ reduction upon mutation of position 344 to Ala^52^.

Extending Replica 2, we further investigated the G_s_ activation mechanism by supervising the opening of the G_s_ alpha-helical domain (AHD), which is considered a necessary step to allow GDP release from the Ras-like domain^53^. We first easily obtained the opening of AHD (**Figure 4b**) and successively supervised the GDP unbinding in a further three replicas, seeded after the AHD opening. In one of these three mwSuMD simulations, the nucleotide dissociated from G_s_ (**Figure 4d**). **Video S7** shows the full G_s_ binding, AHD opening, and GDP release. mwSuMD suggested several structural changes as implicated in GDP dissociation (**Figure 4f-h**): *i*) the AHD opening; *ii*) the conformational change of αG-α4 loop (residues A303^Gα^-P332^Gα^), in concert with a loosening of interactions between D323^Gα^ and K342^6.31^, and *iii*) the rupture of the hydrogen bond between GDP and D295^Gα^ triggered by the movement of αG away from the GDP binding site (**Figure 4f**). The involvement of αG through the αG-α4 during the release of GDP from G_s_ is supported by hydrogen/deuterium exchange experiments^54^, while the role of D275^Gα^ has been probed with functional assays on the G_i_ isoform mutant D272^Gαi^A^55^. Moreover, a similar αG behavior was recently suggested by MD simulations of the G_s_ binding pathway to β_2_AR^56^. We also note that the αG-α4 loop length and aminoacidic composition diverge among G protein isoforms, further suggesting a role in G protein selectivity consistent with the hypothesis that, in different G proteins, distinct domains of the Gα subunit could be responsible for receptor selectivity^57^. However, a more subtle allosteric communication through internal structural elements like the β2-β3 strands, prompted by α5 tilting, could have weakened GDP phosphate binding as previously suggested by other groups^58–60^. Interestingly, no significant conformational changes of the β6-α5 loop happened before or during the GDP dissociation, suggesting that its conformational change as captured in the nucleotide-free GLP-1R:G_s_ complex (**Figure S10**) occurs after the GDP release as a result of the loss of binding stabilization, rather than being an initiator of the GDP dissociation.

## Discussion

Classic MD simulations sample phase space with an efficiency that depends on the energy barrier between neighboring minima. Processes like (un)binding and protein activation require the system to overcome numerous energy barriers, some creating bottlenecks that slow the transition down to the millisecond, or second, time scale. To reduce some of these limits, we have developed, and tested on complex structural events characterizing GPCRs, an energetically unbiased adaptive sampling algorithm, namely multiple walker SuMD, which is based on traditional SuMD, while drawing on parallel multiple replica methods^61,62^, Our simulations propose that remarkable predictivity can be obtained with distance-driven mwSuMD, as demonstrated by the lowest deviation from the experimental AVP:V2R complex. The dissociation of AVP from V2R was simulated much more rapidly by mwSuMD than by SuMD, suggesting it is an efficient tool for studying the dissociation of ligands from GPCRs. This is due to the more extensive sampling obtainable by seeding multiple parallel short simulations instead of a single simulation for batch.

mwSuMD performed similarly to SuMD for the dynamic docking of AVP to V2R when time windows of 600 ps were employed. Time windows of 100 ps remarkably improved mwSuMD. Usually, dynamic docking is performed to either predict the geometry of complexes or sample the binding path of an already-known intermolecular complex, or both. The RMSD of AVP to the experimental coordinates as the supervised metric produced the best results. Consequently, the RMSD should be the metric of choice to study the binding path of well-known intermolecular complexes. The distance, on the other hand, is necessary when limited structural information about the binding mode is available. In the absence of structural information regarding the final bound state, it is possible to sample numerous binding events employing mwSuMD and evaluate the final bound states rank by applying end-point free energy binding methods like the molecular mechanics energies combined with the Poisson– Boltzmann or generalized Born and surface area continuum solvation (MM/PBSA and MM/GBSA^63^) models.

We increased the complexity of binding simulations by considering GLP-1R and the non-peptide agonist PF06882961. Using mwSuMD, we obtained a binding of PF06882961in good agreement with the cryo-EM structure, followed by an active-like conformational transition of GLP-1R. The choice of the metrics supervised was driven by the structural data available^41^ and extensive preparatory MD simulations. However, binding routes are possible from either the bulk solvent or the membrane^36,64,65^. These results show the power of the mwSuMD method, indicating that future applications could include ligand binding from the membrane, or alternative apo receptor conformations to improve the sampling for more difficult receptors.

mwSuMD enabled us to simulate the G_s_ binding to the active GLP-1R and the subsequent GDP release. Our results suggest a concerted effect on GDP binding produced by AHD opening, αG-α4 loop rearrangement, and αG shift away from the GDP site. The full rotation and elongation of α5 as in cryo-EM structures would occur after the GDP release, supporting the role of hidden, metastable interactions as the driving force of G protein coupling and selectivity, as per recent work on GLP-1R^52^.

We stress that a complete understanding of a complex molecular event like G protein coupling requires the collection of numerous G protein binding paths and GDP dissociation events. mwSuMD is designed to yield a mechanistic description of structural events. For this reason, it integrates well with mutagenesis and kinetics experiments. We note that mwSuMD trajectories, since they describe the sequential states along a transition pathway, can represent a precious backbone for further MD sampling aimed at quantifying or predicting the kinetics of the transition. Approaches such as path collective variable metadynamics^66^, Markov State Models^67^, or machine learning models^68,69^ can be informed by mwSuMD. We will address these novel opportunities created by mwSuMD in future work.

In summary, we showcased the applicability domain of mwSuMD to key, but scarcely understood, aspects of GPCR structural biology, pharmacology, and drug design hitherto unaddressed by unbiased simulations. Given the generality and simplicity of its implementation, we anticipate that mwSuMD can be employed to study a wide range of structural phenomena characterizing both membrane and cytosolic proteins. mwSuMD is an ongoing project updated on https://github.com/pipitoludovico/mwSuMD_multiEngine and, in the recent implementation, can exploit ACEMD^70^, NAMD^71^, GROMACS^72^ and OPENMM^73^ as graphic processing units (GPU)-based MD engines while offering the option to run also on CPUs with GROMACS.

## Methods

### Multiple walker SuMD (mwSuMD) protocol

The supervised MD (SuMD) is an adaptive sampling method^74^ based on a tabu-like algorithm for speeding up the simulation of binding events between small molecules (or peptides^75,76^) and proteins^1,19^ without the introduction of any energetic bias. Briefly, during SuMD, a series of short unbiased MD simulations are performed, and after each simulation, the distances between the centers of mass (or the geometrical centers) of the ligand and the predicted binding site (collected at regular time intervals) are fitted to a linear function. If the resulting slope is negative (showing progress towards the target), the next simulation step starts from the last set of coordinates and velocities, otherwise, the simulation is restarted by randomly assigning the atomic velocities.

mwSuMD is designed to increase the sampling from a specific configuration by seeding user-decided parallel replicas (walkers) rather than one short simulation as per SuMD. Since one replica for each batch of walkers is always considered productive, mwSuMD gives more control than SuMD on the total wall-clock time used for a simulation. On the flip side, to maximise mwSuMD, it is optimal to assign one walker per GPU, requiring multiple GPUs to be effective. However, modern multi-threaded GPUs can still employ mwSuMD with a smaller cost in GPU performance. In the implementation for ACEMD used in this work, mwSuMD needs as input the initial coordinates of the system as a pdb file, the coordinates, and the atomic velocities of the system from the equilibration stage, the topology file of the system, and all the necessary force field parameters. The user can decide to supervise one (X) or two metrics (X’, X’’) of the simulated system over short simulations seeded in batches, called walkers. In the former case, either the slope of the linear function interpolating the metric values or a score can be adopted to decide whether to continue the mwSuMD simulation. When the user decides to supervise two metrics, a specific score is used. In the present work, distances between centroids, RMSDs, or the number of atomic contacts between two selections were supervised (**Table S1**). The choice of the metrics is system and problem-dependent, as the RMSD is most useful when the final state is known, while the distance is required when the target state is unknown; details on the scores are given below. The decision to restart or continue mwSuMD after any short simulation is postponed until all the walkers of a batch are collected. The best short simulation is selected and extended by seeding the same number of walkers, with the same duration as the step before.

For each walker, the score for the supervision of a single metric (SMscore) is computed as the square root of the product between the metric value in the last frame (X_last frame_) and the average metric value over the short simulation 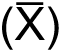:

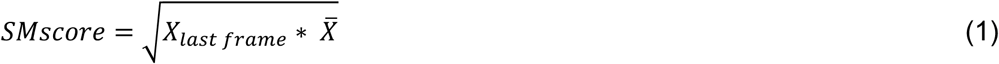

If the metric is set to decrease (e.g. binding or dimerization) the walker with the lowest SMscore is continued, otherwise (e.g. unbinding or outwards opening of domains), the walker with the highest score is continued. Using the SMscore rather than the slope should give more weight to the final state of each short simulation, as it is the starting point for the successive batch of simulations. Considering the average of the metric should favor short simulations consistently evolving in the desired direction along the metric.

If both X’ and X’’ are set to increase during the mwSuMD simulations, the score for the supervision of two metrics (DMscore) on each walker is computed as follows:

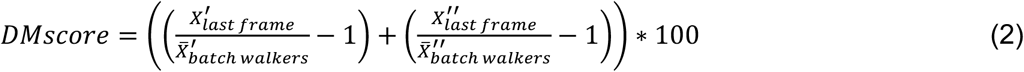

Where X’_last frame_ and X’’_last frame_ are the metrics values in the last frame, while 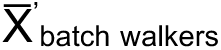 and 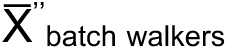 represent the average value of the two metrics over all the walkers in the batch. Subtracting the value 1 to the metric ratio ensures that if one of the two metrics from the last frame (X’_last frame_ or X^’’^_last frame_) is equal to the average 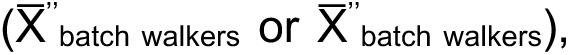 then that metric addend is null, and DMscore depends only on the remaining metric. If any of the two metrics is set to decrease, then the corresponding component in Equation 2 is multiplied by - 1 to maintain a positive score. Considering the average value of the two metrics over all the walkers rather than only over the considered walker should be more representative of the system evolution along the defined metric. In other words, the information about the metric is taken from all the walkers to better describe the evolution of the system.

The DMScore is designed to preserve some degree of independence between the two metrics supervised. Indeed, if the variation of one of them slows down and gets close to zero, the other metric is still able to drive the system’s evolution. It should be noted that DMScore works at its best if the two metrics have similar variations over time, as in the case of distance and RMSD (both of which are distance-based). Notably, differently from SuMD, when a walker is extended by seeding a new batch of short simulations and the remaining walkers are stopped, the atomic velocities are not reassigned. This allows the simulations to be as short as a few picoseconds if desired, without introducing artifacts due to the thermostat latency to reach the target temperature (usually up to 10-20 ps when a simulation is restarted, reassigning the velocities of the atoms).

The current implementation of mwSuMD is for Python3 and exploits MDAnalysis^77^ and MDTRaj^78^ modules.

### Force field, ligands parameters, and general systems preparation

The CHARMM36^79,80^/CGenFF 3.0.1^81–83^ force field combination was employed in this work. Initial ligand force field, topology and parameter files were obtained from the ParamChem webserver^81^. Restrained electrostatic potential (RESP)^84^ partial charges were assigned to all the non-peptidic small molecules but adrenaline and guanosine-5’-diphosphate (GDP) using Gaussian09 (HF/6-31G* level of theory) and AmberTools20.

Six systems were prepared for MD (**Table S1**). Hydrogen atoms were added using the pdb2pqr^85^ and propka^86^ software (considering a simulated pH of 7.0); the protonation of titratable side chains was checked by visual inspection. The resulting receptors were separately inserted in a 1-palmitoyl-2-oleyl-sn-glycerol-3-phosphocholine (POPC) bilayer (previously built by using the VMD Membrane Builder plugin 1.1, Membrane Plugin, Version 1.1. at: http://www.ks.uiuc.edu/Research/vmd/plugins/membrane/), through an insertion method^87^. Receptor orientation was obtained by superposing the coordinates on the corresponding structure retrieved from the OPM database^88^. Lipids overlapping the receptor transmembrane helical bundle were removed and TIP3P water molecules^89^ were added to the simulation box employing the VMD Solvate plugin 1.5 (Solvate Plugin, Version 1.5. at <http://www.ks.uiuc.edu/Research/vmd/plugins/solvate/). Finally, overall charge neutrality was reached by adding Na^+^/Cl^-^ counter ions up to the final concentration of 0.150 M), using the VMD Autoionize plugin 1.3 (Autoionize Plugin, Version 1.3. at <http://www.ks.uiuc.edu/Research/vmd/plugins/autoionize/).

### System equilibration and general MD settings

The MD engine ACEMD3^70^ was employed for both the equilibration and productive simulations. The equilibration was achieved in isothermal-isobaric conditions (NPT) using the Berendsen barostat^90^ (target pressure 1 atm) and the Langevin thermostat^91^ (target temperature 300 K) with low damping of 1 ps^-1^. For the equilibration (integration time step of 2 fs): first, clashes between protein and lipid atoms were reduced through 1500 conjugate-gradient minimization steps, then a positional constraint of 1 kcal mol^-1^ Å^-2^ on all heavy atoms was gradually released over different time windows: 2 ns for lipid phosphorus atoms, 60 ns for protein atoms other than alpha carbon atoms, 80 ns for alpha carbon atoms; a further 20 ns of equilibration was performed without any positional constraints.

Productive trajectories (**Table S1**) were computed with an integration time step of 4 fs in the canonical ensemble (NVT). The target temperature was set at 300 K, using a thermostat damping of 0.1 ps^-1^; the M-SHAKE algorithm^92,93^ was employed to constrain the bond lengths involving hydrogen atoms. The cut-off distance for electrostatic interactions was set at 9 Å, with a switching function applied beyond 7.5 Å. Long-range Coulomb interactions were handled using the particle mesh Ewald summation method (PME)^94^ by setting the mesh spacing to 1.0 Å.

### Vasopressin binding simulations

The vasopressin 2 receptor (V_2_R) in complex with vasopressin (AVP) and the G_s_ protein^95^ was retrieved from the Protein Data Bank^96^ (PDB 7DW9). The G_s_ was removed from the system and the missing residues on ECL2 (G185-G189) were modeled from scratch using Modeller 9.19^97^, considering the solution with the lowest DOPE score out of ten conformations produced. AVP was placed away from V_2_R in the extracellular bulk and the resulting system was prepared for MD simulations and equilibrated as reported above.

During SuMD simulations, the distance between the centroids of AVP residues C1-Q4 (backbone and side chains), anticipated to bind deep into V_2_R, and the V_2_R residues lining the peptide binding site Q96, Q174, Q291, and L312 (Cα atoms only) was supervised over time windows of 600 ps or 100 ps (**Table S1**).mwSuMD simulations considered the same distance, the RMSD of AVP residues C1-Q4 to the experimental bound complex or the combination of the two during time windows of 600 ps (3 walkers) or 100 ps (10 walkers) (**Table S1**). Slope, SMscore, or DMscore (see Methods section **mwSuMD protocol**) was used in the different mwSuMD replicas performed (**Table S1**). Simulations were stopped after 300 ns (time window duration = 600 ps) or 50 ns (time window duration = 100 ps) of total SuMD or mwSuMD simulation time.

### Vasopressin unbinding simulations

The V_2_R:AVP complex was prepared for MD simulations and equilibrated as reported above. During both SuMD and mwSuMD simulations (**Table S1**), the distance between the centroids of AVP residues C1-Q4 (backbone and side chains) and V_2_R residues Q96, Q174, Q291, and L312 (Cα atoms only) was supervised over time windows of 100 ps (10 walkers seeded for mwSuMD simulations). Replicas were stopped when the AVP-V_2_R distance reached 40 Å.

### GLP-1R:PF06882961 binding simulations

The inactive glucagon-like peptide receptor (GLP-1R) was retrieved from the Protein Data Bank^96^ (PDB 6LN2)^98^. Fab and the intracellular negative allosteric modulator were removed, and missing residues in the stalk (129-134) and ICL2 (256 to 263) were modeled with Modeller 9.19, considering the solutions with the lowest DOPE score out of ten conformations produced. The PF06882961 initial conformation was extracted from the complex with the fully active GLP-1R^45^ (PDB 7LCJ) and placed away from GLP-1R in the extracellular bulk. The resulting system was prepared for MD simulations and equilibrated as reported above. CGenFF dihedral force field parameters of PF06882961 with the highest penalties (dihedrals NG2R51-CG321-CG3C41-CG3C41 (penalty=143.5) and NG2R51-CG321-CG3C41-OG3C51 (penalty=152.4)) were optimized (**Figure S11**) employing Gaussian09 (geometric optimization and dihedral scan at HF/6-31g(d) level of theory) and the VMD force field toolkit plugin^99^.

Four classic MD replicas, for a total of 8 μs, were performed on the inactive receptor (prepared for MD simulations and equilibrated as reported above) to assess the possible binding path to the receptor TMD and therefore decide the initial position of PF06882961 in the extracellular bulk of the simulation box. A visual inspection of the trajectories suggested three major conformational changes that could allow ligand access to the TMD (**Figure S12**). Transitory openings of the ECD (distance Q47^ECD^ - S310^ECL2^), TM6-TM7 (distance H363^6.52^ - F390^7.45^), and TM1-ECL1 (distance E138^1.33^ and W214^ECL1^) were observed. Since the opening of TM1-ECL1 was observed in two replicas out of four, we placed the ligand in a favorable position for crossing that region of GLP-1R.

mwSuMD simulations (**Table S1**) were performed stepwise to dock the ligand within GLP-1R first and then relax the receptor towards the active state. The PF06882961 binding was obtained by supervising at the same time the distance between the ligand’s heavy atoms centroid and the centroid of GLP-1R TM7 residues L379^7.34^-F381^7.36^ (Cα atoms only), which are part of the orthosteric site, and the RMSD of the ECD (residues W33^ECD^-W120^ECD^, Cα atoms only) to the active state (PDB 7LCJ) until the former distance reached 4 Å. In the second phase of mwSuMD, the RMSD of the ECD (residues W33^ECD^-W120^ECD^, Cα atoms only) and the ECL1 to the active state (PDB 7LCJ, Cα atoms of residues M204^2.74^-L224^3.27^) were supervised until the latter reached less than 4 Å. During the third phase, the RMSD of PF06882961, as well as the RMSD of ECL3 (residues A368^6.57^-T378^7.33^, Cα atoms), were supervised until the former reached values lower than 3 Å. In the last mwSuMD step, only the RMSD of TM6 (residues I345^6.34^-F367^6.56^, Cα atoms) to the active state (PDB 7LCJ) was supervised until less than 5 Å. RMSDs were computed after superimposition on TM2, ECL1, and TM3 residues 170-240 (Cα atoms), which is the GLP-1R less flexible part^50^.

### Membrane-anchored G_s_ protein:GLP-1R simulations and GDP dissociation

The PDB 6EG8 was processed through Charmm-GUI^100^ to palmitoylate residue C3^Gαi^ and geranylgeranylate residue C65^Gγ^. The resulting system was inserted into a 120 x 120 Å POPC membrane and previously built by using the VMD Membrane Builder plugin 1.1, Membrane Plugin, Version 1.1. at: http://www.ks.uiuc.edu/Research/vmd/plugins/membrane. Lipids overlapping the palmitoyl and geranylgeranyl groups were removed and TIP3P water molecules^89^ were added to the simulation box by means of the VMD Solvate plugin 1.5 (Solvate Plugin, Version 1.5. at <http://www.ks.uiuc.edu/Research/vmd/plugins/solvate/).

Finally, overall charge neutrality was reached by adding Na^+^/Cl^-^ counter ions up to the final concentration of 0.150 M), using the VMD Autoionize plugin 1.3 (Autoionize Plugin, Version 1.3. at <http://www.ks.uiuc.edu/Research/vmd/plugins/autoionize/). The first stage of equilibration was performed as reported above (Methods section **System equilibration and general MD settings**) for 120 ns, followed by a second stage in the NVT ensemble for a further 1 μs without any restraints to allow the membrane-anchored heterotrimeric G_s_ protein to stabilize within the intracellular side of the simulation box. After this two-stage long equilibration, GLP-1R from the final frame of the activation simulation (in complex with PF06882961) was manually inserted into the equilibrated membrane above the G_s_ protein using the corresponding structure retrieved from the OPM database as a reference, and the system further equilibrated for 120 ns as reported above (Methods section **System equilibration and general MD settings**). The GLP-1R-G_s_ system was then subjected to three simulations (**Table S1**). Each mwSuMD replica was interrupted by 500 ns of classic MD twice, to relax the system during the transition. In the supervised stages, the distance between residues M386-L394 ^Gαs^ (all-atoms centroid) of helix 5 (α5) and the GLP-1R intracellular residues R176^2.46^, R348^6.37^, S352^6.41^, and N405^7.60^ (Cα atoms only) was monitored, seeding three walkers of 200 ps each.

The AHD opening was simulated starting from the GLP-1R:G_s_ binding mwSuMD replica with the final lowest G_s_ RMSD, the lowest PF06882961 binding energy and the highest GDP binding energy (Replica 2 in **Figure 4e**) by supervising the distance between AHD residues G70-R199 ^Gαs^ and K300-L394^Gαs^ (all-atoms centroids) during three walkers of 100 ps each. 300 ns of classic MD was performed to relax the system. Finally, the GDP unbinding was supervised as the distance between GDP (all-atoms centroid) and residues E50^Gαs^, K52^Gαs^, T55^Gαs^, K293^Gαs^, and V367^Gαs^ (Cα atoms only) of G_αs_; five walkers were used in a 50 ps long mwSuMD simulations.

### MD Analysis

Interatomic distances were computed through MDAnalysis^77^; root mean square deviations (RMSD) were computed using VMD^101^ and MDAnalysis^77^. Interatomic contacts and ligand-protein hydrogen bonds were detected using the GetContacts scripts tool (https://getcontacts.github.io), setting a hydrogen bond donor-acceptor distance of 3.3 Å and an angle value of 120° as geometrical cut-offs. Contacts and hydrogen bond persistency are quantified as the percentage of frames (over all the frames obtained by merging the different replicas) in which protein residues formed contacts or hydrogen bonds with the ligand.

The MMPBSA.py^102^ script, from the AmberTools20 suite (The Amber Molecular Dynamics Package, at http://ambermd.org/), was used to compute molecular mechanics energies combined with the generalized Born and surface area continuum solvation (MM/GBSA) method or the molecular mechanics Poisson-Boltzmann surface area (MM/PBSA) approach, after transforming the CHARMM psf topology files to an Amber prmtop format using ParmEd (documentation at <http://parmed.github.io/ParmEd/html/index.html).

Supplementary Videos were produced employing VMD and avconv (at https://libav.org/avconv.html). Molecular graphics images were produced using the UCSF Chimera^103^ (v1.14).

### Numbering system

Throughout the manuscript, the Ballesteros-Weinstein residues numbering system for class A^104^ and the Wootten residues numbering system for class B GPCRs^105^ are adopted.

## ASSOCIATED CONTENT

Supporting Information (PDF) Supporting Videos S1-S7 (mpg).

All the MD trajectories (stripped of POPC, water molecules, and ions) and topology files (psf and pdb) are available here: https://zenodo.org/record/7944479

The mwSuMD software version used in this study is available at: https://github.com/pipitoludovico/mwSuMD

## Author Contributions

GD and CAR conceived and supervised the project; GD designed and implemented the software and planned the simulations; GD, LP, RMR, TW, and PG carried out the simulations; GD analyzed the data; GD, AC, SM, and CAR interpreted the results, GD wrote the manuscript with input from CAR, AC, and SM; all the authors edited and reviewed the final version of the manuscript.

## Acknowledgments

GD is a member of the GPCRs-focused European COST action ERNEST. CAR is grateful for a Royal Society Industry Fellowship. GD and CAR are grateful for support from the BBSRC (BB/W016974/1) and Diabetes UK (BDA 20/0006307).

