## Supplementary for "Hidden GPCR structural transitions addressed by multiple walker supervised molecular dynamics (mwSuMD)"

### Supplementary Videos

**Video S1. AVP binding (dynamic docking) to $V_2R$.** Two-view of the mwSuMD replica better reproducing the AVP:$V_2R$ experimental complex. Left side: $V_2R$ is represented in white quick surface, while AVP is in transparent quick surface and green stick; right side: $V_2R$ is represented in white ribbon and cyan stick, while AVP is a green stick. The bound AVP conformation from PDB 7DW9 is reported as a reference in transparent orange ribbon and stick.

**Video S2. AVP unbinding from $V_2R$.** Two-view of the five mwSuMD replicas performed. Left side: $V_2R$ is represented in white quick surface, while AVP is in transparent quick surface and green stick; right side: $V_2R$ is represented in white ribbon and cyan stick, while AVP is a green stick.

**Video S3. $G_s$ binding to $\beta_2AR$.** The inactive $G_s$ ($G_\alpha$ subunit in orange, $G_\beta$ subunit in green, and $G_\gamma$ subunit in yellow) recognizes $\beta_2AR$ (black ribbon) bound to epinephrine (van der Waals spheres). The experimental $\beta_2AR$:$G_s$ cryo-EM complex is reported in white ribbon for reference.

**Video S4. $G_i$ binding to $A_1R$.** The $G_i$ ($G_\alpha$ subunit in magenta, $G_\beta$ subunit in green, and $G_\gamma$ subunit in yellow) recognizes $A_1R$ (blue ribbon) bound to adenosine (not shown). The experimental $A_1R$:$G_i$ cryo-EM complex is reported in transparent ribbon for reference.

**Video S5. Morph of the PF06882961 binding and GLP-1R activation.** The first and last frames of the PF06882961 binding simulations have been interpolated to produce a smoothed representative transition.

**Video S6. PF06882961 binding and GLP-1R activation.** PF06882961 is represented in cyan van der Waals spheres, while GLP-1R is orange ribbon; the GLP-1R experimental active conformation is reported in transparent green ribbon as reference.

**Video S7. Gs binding to GLP-1R and GDP release.** The Gs protein binds GLP-1R (light grey) in complex with PF06882961 (stick representation) before the alpha-helical domain opens and allows GDP release from $G_{s\alpha}$. The $G_{s\alpha}$ subunit is yellow, $G_\beta$ is dark gray, and $G\gamma$ is orange.

**Supplementary Methods**

**$G_s$ protein:$\beta_2$ AR binding simulations**

mwSuMD simulations started from the intermediate, agonist-bound conformation of the $\beta_2$-AR and the inactive $G_s$ to resemble pre-coupling conditions. The full-length model of the adrenergic $\beta_2$ receptor ($\beta_2$ AR) in an intermediate active state was downloaded from GPCRdb (https://gpcrdb.org/). The full agonist adrenaline (ALE) was inserted in the orthosteric site by superposition with the PDB ID 4LDO (fully active $\beta_2$ AR)[1]. The structure of the inactive, GDP-bound $G_s$ protein[2] was retrieved from the Protein Data Bank[3] (PDB ID 6EG8) and placed in the intracellular bulk. The resulting system ($G_s$ > 50 Å away from ($\beta_2$ AR) was prepared for MD simulations and equilibrated as reported above. The PDB ID 3SN6 (fully active $\beta_2$ AR in complex with $G_s$[4]) was used as the reference for RMSD computations. Three mwSuMD replicas (**Table S1**) were performed supervising at the same time the distance between the helix 5 (α5) $G_\alpha$s residues R385-L395 (Cα atoms centroid) and the $\beta_2$ AR (residues V31-P330 Cα atoms centroid) as well as the RMSD (superimposing on $\beta_2$ AR residues 70-170 Cα atoms) of $\beta_2$ AR TM6 residues C265-I278 (Cα atoms only) to the fully active state, during 100 ps time windows (5 walkers). To monitor the progression of the simulations, we computed the RMSD of the Cα atoms of the Gα (residues 11 to 43 and resid 205 to 394) and Gβ subunits (residues 3 to 340) to the experimental complex[4] (**Video S3, Figure S9a,b**). The flexibility of $G_s\beta$ is backed by both MD and cryo-EM data suggesting G protein rocking motions around $G_s\alpha$:receptor interactions[5,6].

**Membrane-anchored Gi protein:$A_1$R simulations**

Since the full-length structure of the inactive human Gi protein has not been yet resolved by X-ray or cryo-EM, it was modeled by superimposing the AlphaFold2[7] AI models of the Gαi (P63096-F1), Gβ (Q9HAV0-F1), and Gγ (P50151-F1) subunits to the PDB file 6EG8 (a $G_s$ heterotrimer). The resulting homotrimer (without GDP) was processed through Charmm-GUI[8] to palmitoylate residue $C3^{G\alpha i}$ and geranylgeranylate residue $C65^{G\gamma}$ [9,10]. The side chains of these two lipidated residues were manually inserted into a 120 x 120 Å POPC membrane and previously built by using the VMD Membrane Builder plugin 1.1, Membrane Plugin, Version 1.1. at: http://www.ks.uiuc.edu/Research/vmd/plugins/membrane/. Lipids overlapping the palmitoyl and geranylgeranyl groups were removed and TIP3P water molecules[11] were added to the simulation box by means of the VMD Solvate plugin 1.5 (Solvate Plugin, Version 1.5. at <http://www.ks.uiuc.edu/Research/vmd/plugins/solvate/). Finally, overall charge neutrality was reached by adding $Na^+/Cl^-$ counter ions up to the final concentration of 0.150 M), using the VMD Autoionize plugin 1.3 (Autoionize Plugin, Version 1.3. at <http://www.ks.uiuc.edu/Research/vmd/plugins/autoionize/). The first stage of equilibration was performed as reported above (Methods section **System equilibration and general MD settings**) for 120 ns, followed by a second stage in the NVT ensemble for a further 1 μs without any restraints to allow the membrane-anchored heterotrimeric Gi protein to stabilize within the intracellular side of the simulation box. After this two-stage long equilibration, the active state $A_1$R in complex with adenosine (PDB 6D9H) was manually inserted into the equilibrated membrane above the Gi protein using the corresponding structure retrieved from the OPM database as a reference, and the system further equilibrated for 120 ns as reported in the Methods section **System equilibration and general MD settings**. The $A_1$R-Gi system was

then subjected to a 1 μs-long classic MD simulation and a mwSuMD simulation (**Table S1**). During the mwSuMD simulation, the RMSD (superimposing on $A_1R$ residues 40-140 Cα atoms) of helix 5 (α5) $G_{\alpha i}$ residues 329-354 to the PDB 6D9H was supervised, seeding three walkers of 100 ps each until the productive simulation time reached 50 ns (total simulation time 150 ns).

### Supplementary Figures

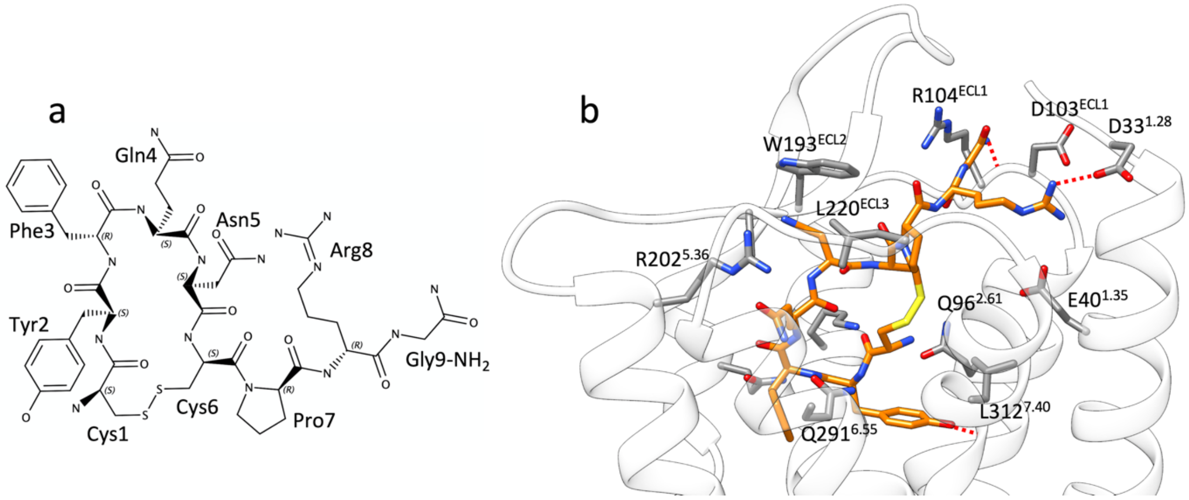


**Figure S1.** Arginine vasopressin (AVP) nonapeptide. **a)** chemical structure and **b**) binding more within the $V_2R$ orthosteric binding site; AVP is represented in orange stick, while $V_2R$ is in white ribbon and grey stick.

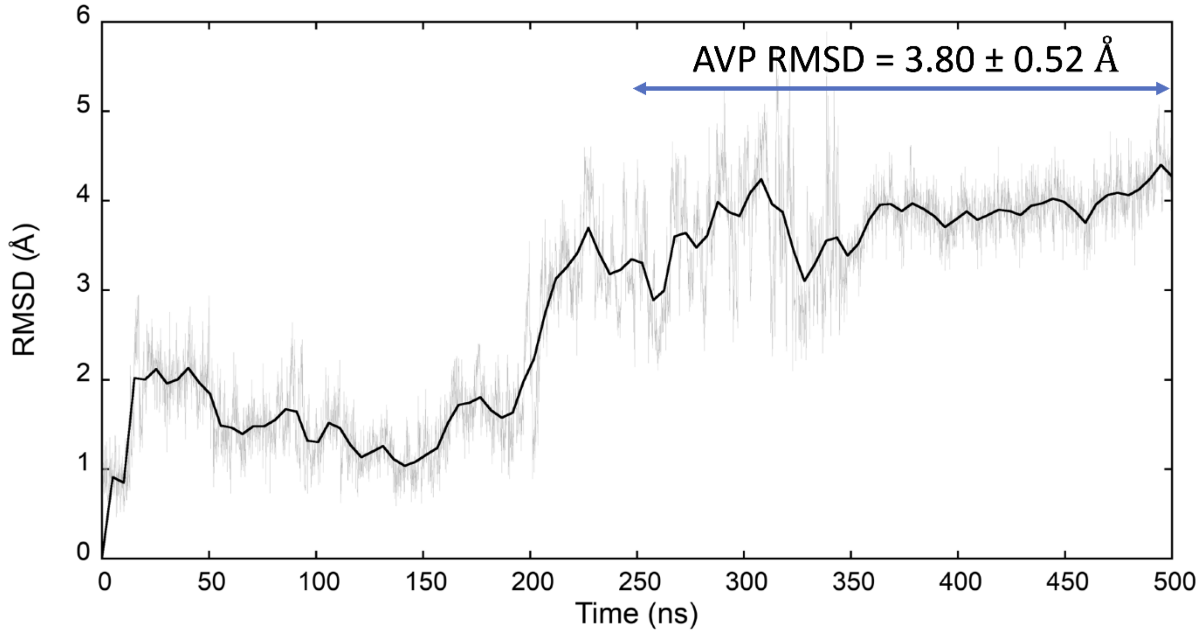


**Figure S2.** Classic MD simulation of the AVP:$V_2R$ complex (PDB ID 7DW9)**.** The average AVP RMSD (computed on the Cα atoms) after stabilization of the system (second half of the trajectory) is indicated.

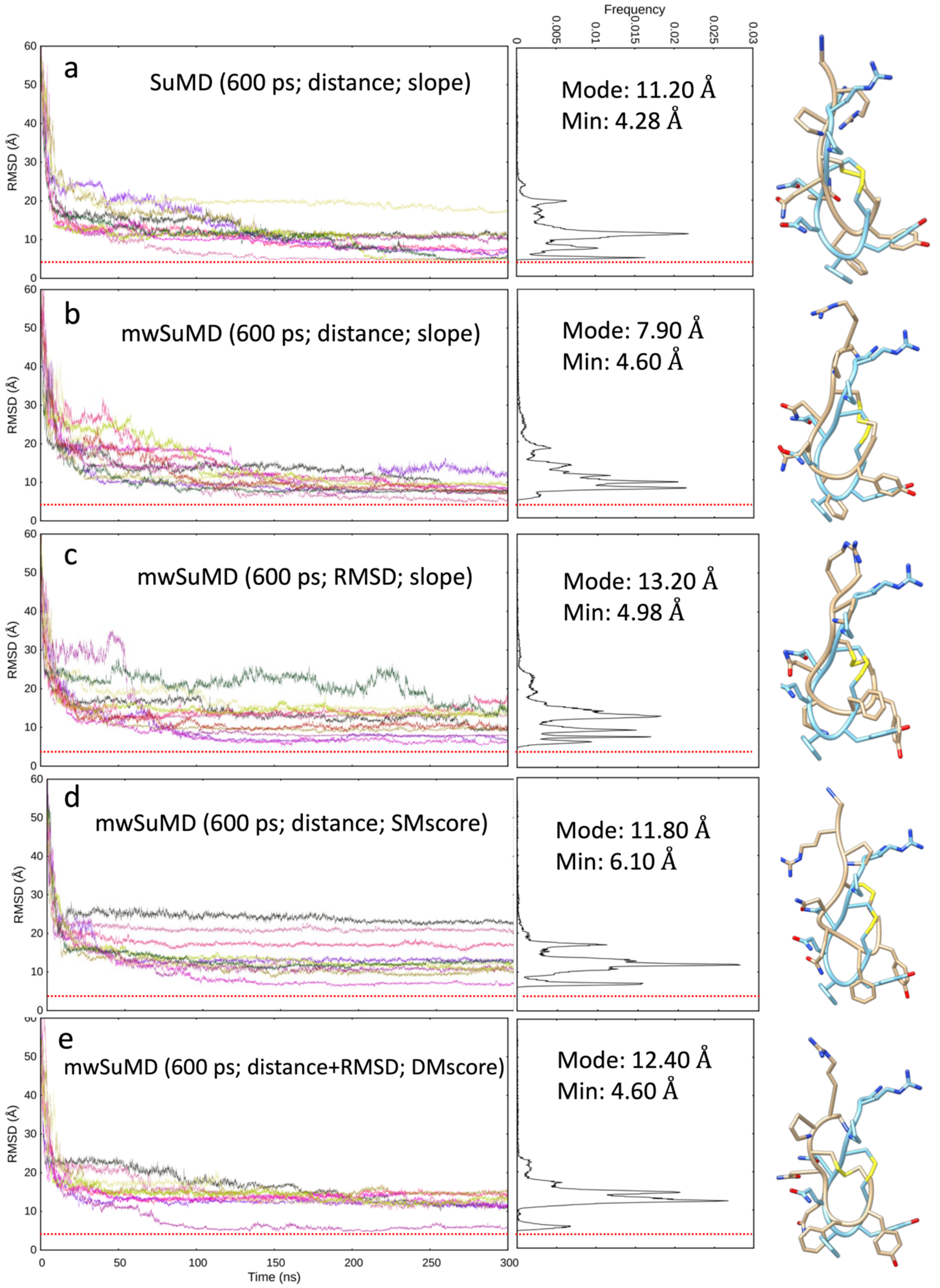


**Figure S3.** AVP SuMD and mwSuMD binding simulations to $V_2R$ (600 ps time windows). For each set of settings **(a-e),** the RMSD of AVP C$\alpha$ atoms to the cryo-EM structure 7DW9 is reported during the time course of each SuMD (**a**) or mwSuMD (**b-e**) replica, alongside the RMSD values distribution and the snapshot corresponding to the lowest RMSD (AVP from the cryo-EM structure 7DW9 is in a cyan stick representation, while AVP from simulations is in a tan stick representation). A complete description of the simulation settings is reported in Table S1 and the Methods section. The dashed red line indicates the AVP RMSD during a classic (unsupervised) equilibrium MD simulation of the X-ray AVP:$V_2$R complex (**Figure S2**).

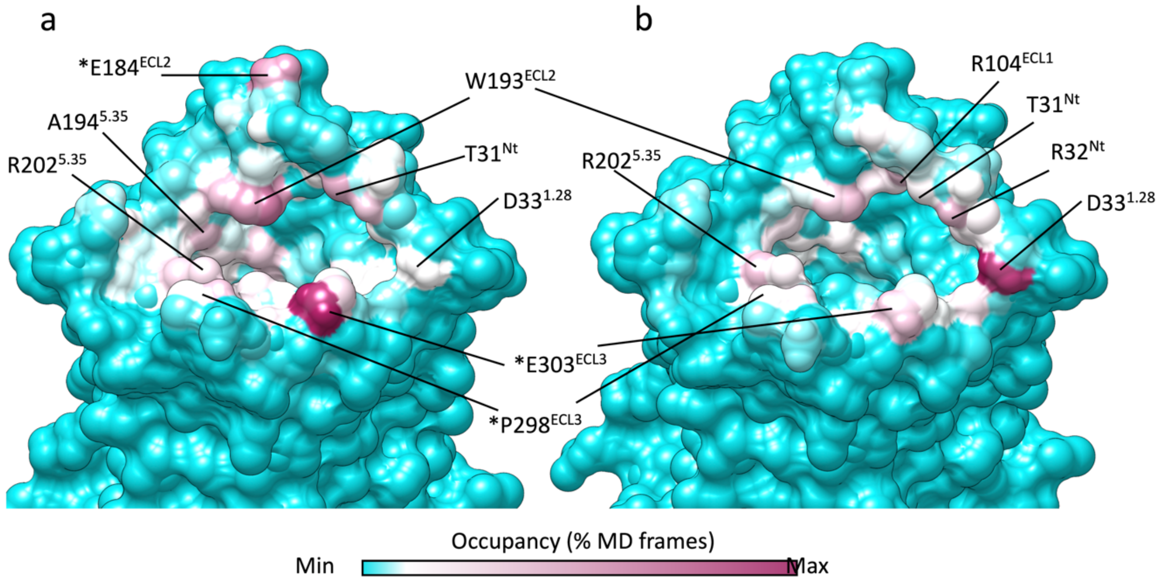


**Figure S4.** $V_2$R residues involved in mwSuMD simulations of AVP. **a)** Binding (dynamic docking); **b**) unbinding simulations.

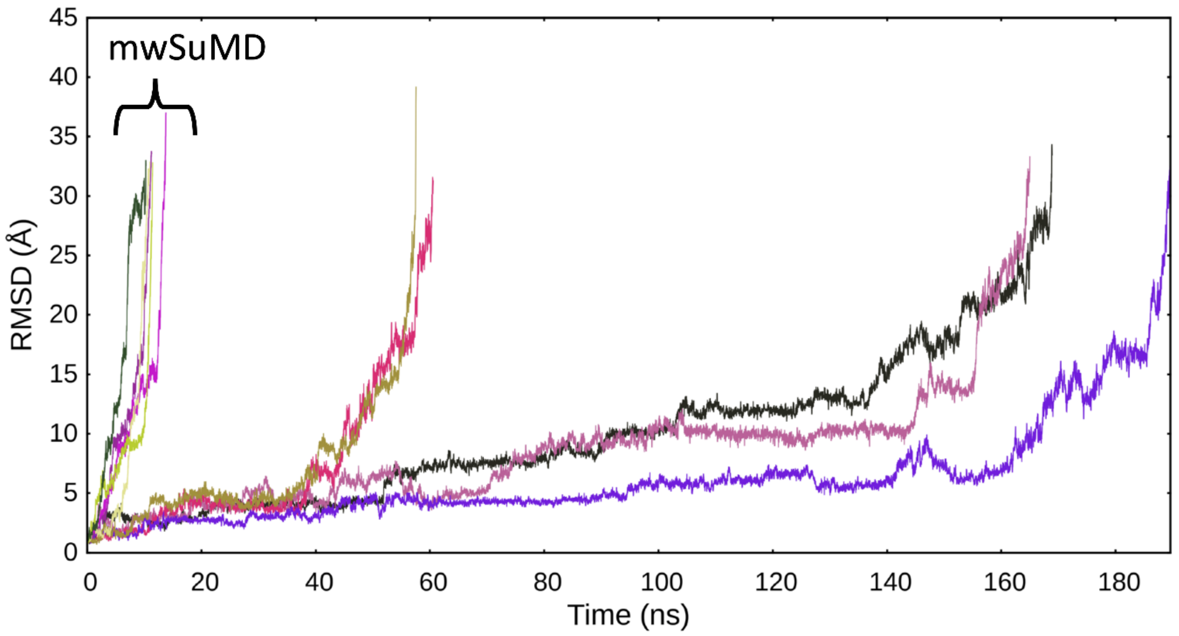


**Figure S5.** Dissociation of AVP from $V_2$R. The RMSD of AVP to the initial bound state is reported during the time course of five replicas of SuMD and mwSuMD, respectively.

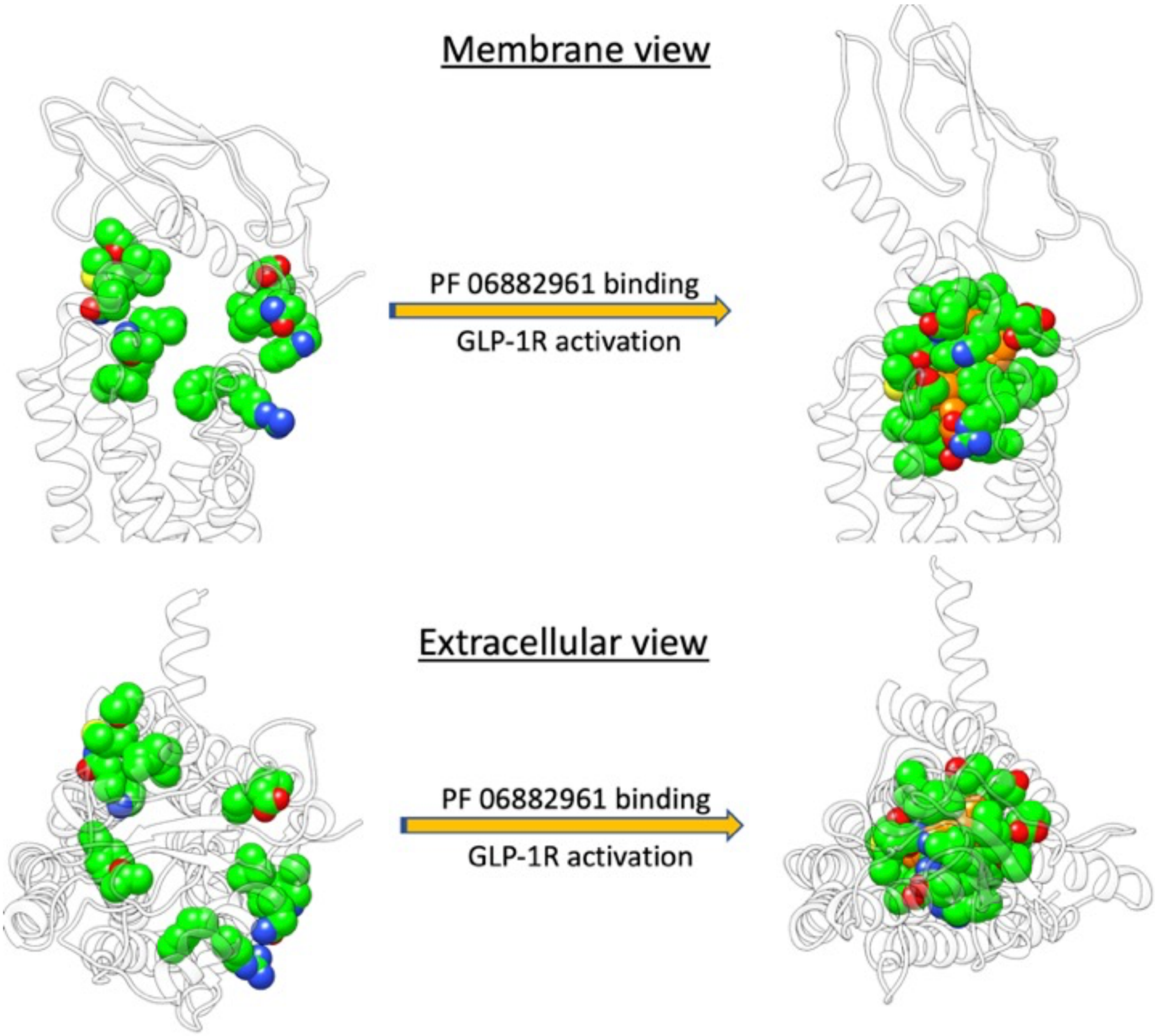


**Figure S6.** PF06882961 binding site in the apo and holo GLP-1R**.** In the apo GLP-1R (left transparent ribbon) the residues forming the binding site of the NPA PF06882961 are scattered due to the different conformation of the receptor.

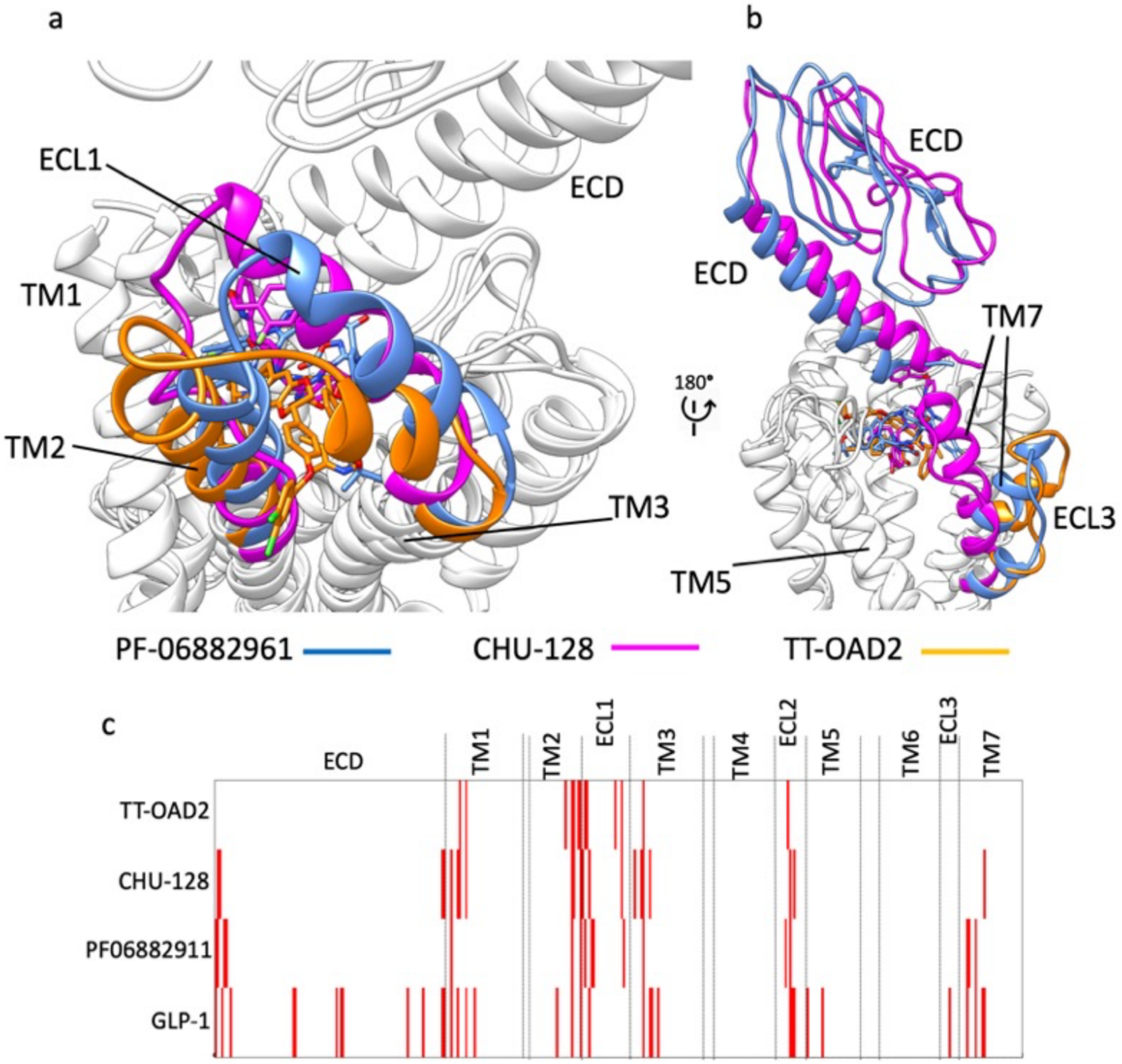


**Figure S7.** NPAs stabilize divergent GLP-1R active conformations**. a)** Different orientations of TM2, ECL1, and TM3; **b**) divergent conformations of ECD, ECL3 and TM7; **c**) interaction fingerprint of NPAs (TT-OAD2, CHU-128, and PF06882911) and the endogenous agonist peptide GLP-1. Red bars indicate interactions with distinct residues in the GLP-1R.

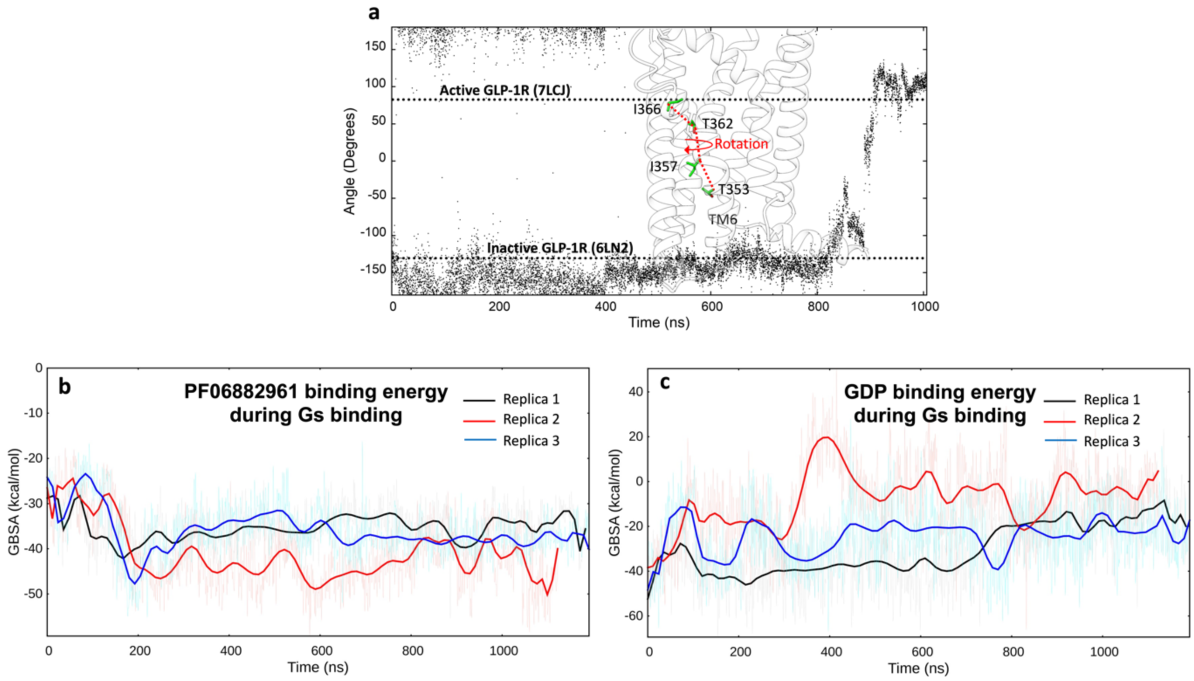


**Figure S8. a)** Rotation of TM6 during GLP-1R activation simulated by mwSuMD. The angle was measured as the dihedral formed by the backbone alpha carbon atoms T353, I357, T362, and I366. The angle values of the inactive and active cryo-EM structures are reported as references; **b**) PF06882961 MM-GBSA binding energy during $G_s$ binding; **d**) GDP MM-GBSA binding energy during $G_s$ binding;

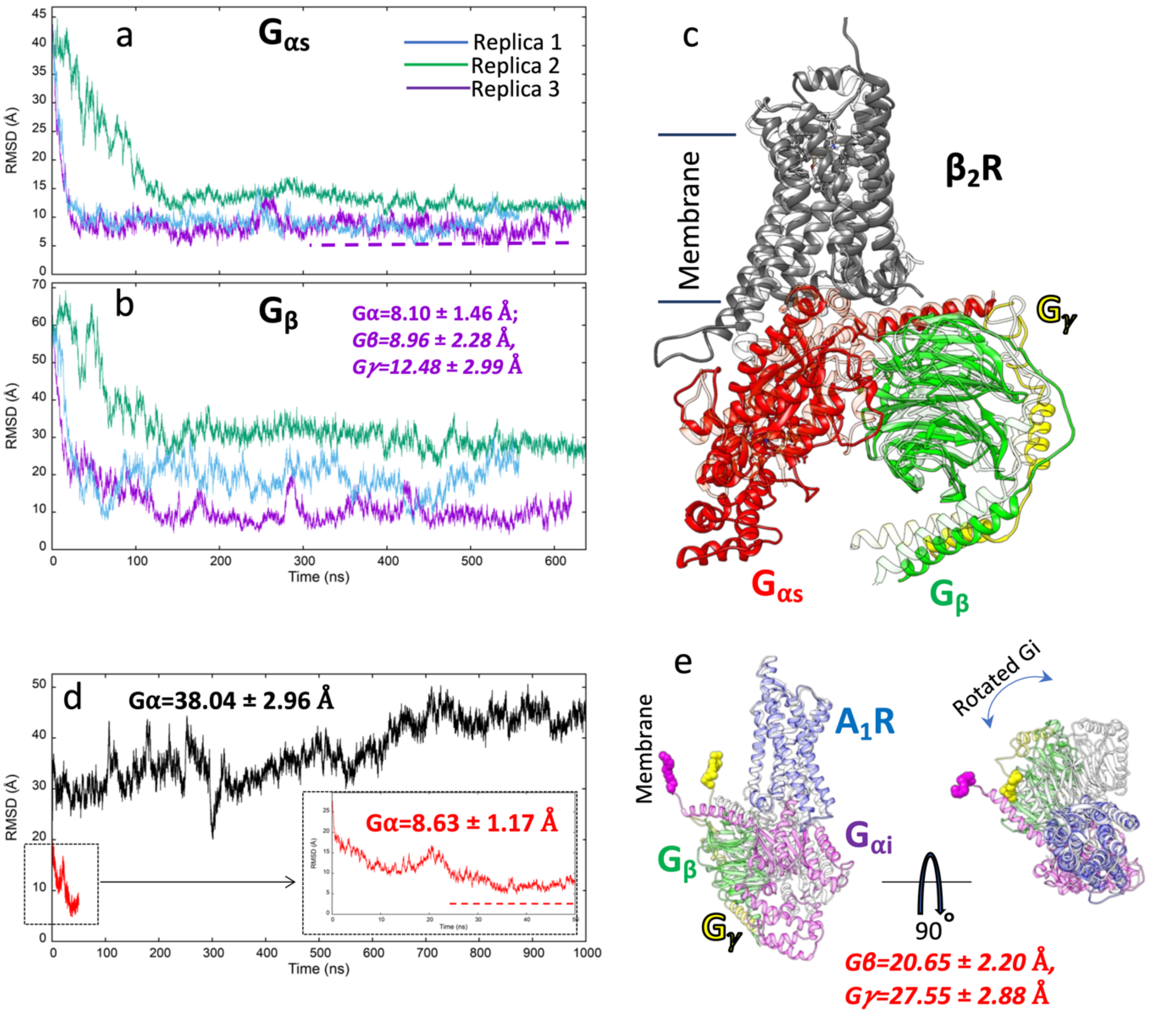


**Figure S9. G protein binding simulations to $\beta_2$AR and $A_1$R. a)** RMSD of $G_s\alpha$ to the experimental complex (PDB 3NS6) during three mwSuMD replicas; **b)** RMSD of $G_s\beta$ to the experimental complex (PDB 3NS6) during three mwSuMD replicas; **c)** superposition of the experimental $G_s$:$\beta_2$ AR complex (transparent ribbon) and the MD frame with the lowest $G_s\alpha$ RMSD **(**3.94 Å**); d**) RMSD of $G_i\alpha$ (residues 243-355) to the $A_1$R experimental complex (PDB 6D9H) during a mwSuMD simulation (red, magnified in the box) and a 1000-ns long classic MD simulation (black); **e**) two-view superposition of the experimental $G_i$:$A_1$ R complex (transparent ribbon) and the MD frame with the lowest $G_i\alpha$ RMSD **(**4.82 Å**).**

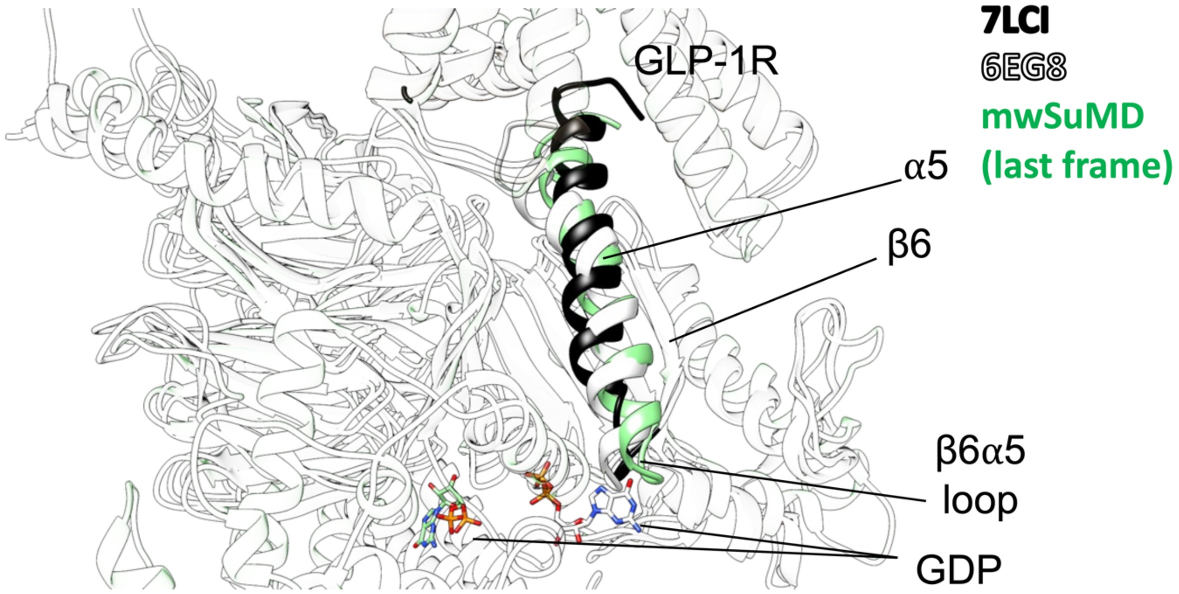


**Figure S10**. Comparison of the inactive (6EG8, white), nucleotide-free (7LCI, black), and GDP-dissociated (mwSuMD simulation, green) β6-α5 loop. Bound (white, 6EG8) and dissociated (green, mwSuMD) GDP is shown as a stick representation.

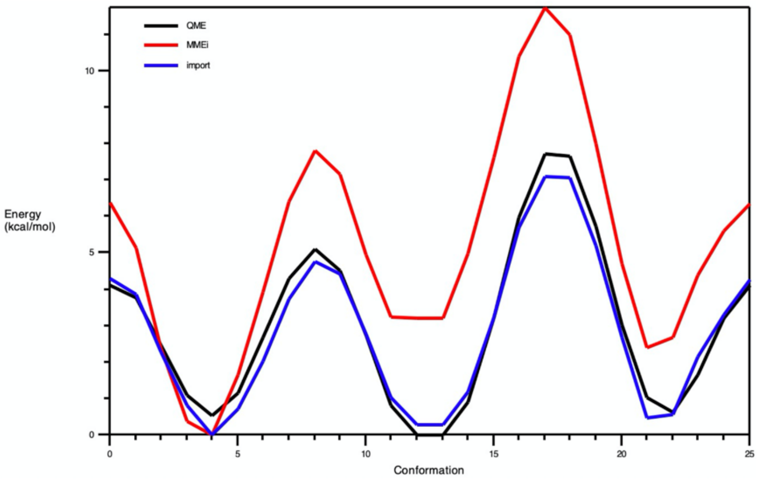


**Figure S11**. Potential energy surface derived from the scan of the PF06882961 rotatable bond through which the dihedrals NG2R51-CG321-CG3C41-CG3C41 (penalty=143.5) and NG2R51-CG321-CG3C41-OG3C51 (penalty=152.4) pass. The curve obtained from the original CGenFF parameters (red) was optimized (blue) to resemble the energy profile from quantum-mechanics computations at the HF/6-31g(d) level of theory (black).

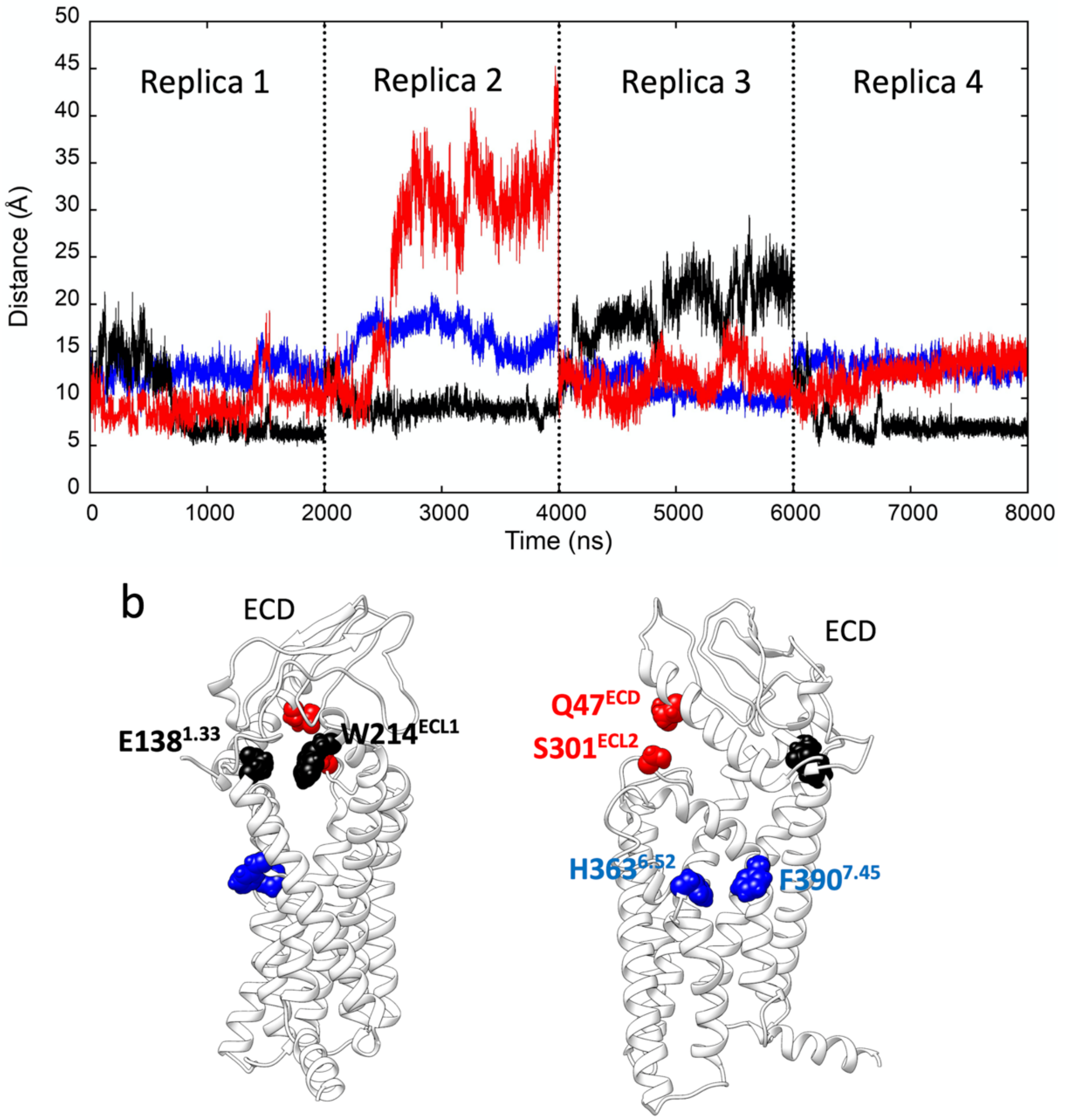


**Figure S12. GLP.1R conformational changes during 8 μs of classic MD. a)** Distances between key residues E138$^{1.33}$ and W214$^{ECL1}$ (black), H363$^{6.52}$ and F390$^{7.45}$ (blue) and Q47$^{ECD}$ and S310$^{ECL2}$ (red) during the course of the simulation. **(b)** The position of the three couples of residues is reported as two-side views.

### Supplementary Tables

**Table S1. Summary of all the simulations performed and the settings employed.**

| System | # Replicas | TW Duration | # Walkers | Metric | Acceptance |
|---|---|---|---|---|---|
| **$V_2R$:AVP Complex** | 1 classic MD | 500 ns | N/A | N/A | N/A |
| **$V_2R$:AVP** | 9 (SuMD) | 600 ps | N/A | Distance | Slope |
| | 10 (mwSuMD) | 600 ps | 3 | Distance | Slope |
| | 11 (mwSuMD) | 600 ps | 3 | RMSD | Slope |
| | 8 (mwSuMD) | 600 ps | 3 | Distance | SMscore |
| | 9 (mwSuMD) | 600 ps | 3 | Distance and RMSD | DMscore |

| | | | | | |
|---|---|---|---|---|---|
| **Binding** | 12 (mwSuMD) | 100 ps | 10 | Distance and RMSD | DMscore |
| | 11 (mwSuMD) | 100 ps | 10 | Distance | SMscore |
| | 11 (mwSuMD) | 100 ps | 10 | RMSD | SMscore |
| | 10 (SuMD) | 100 ps | N/A | Distance | Slope |
| **$V_2$R:AVP Unbinding** | 5 (SuMD) | 100 ps | N/A | Distance | Slope |
| | 5 (mwSuMD) | 100 ps | 10 | Distance | SMscore |
| **$\beta_2$-AR:Gs protein Binding** | 3 (mwSuMD) | 100 ps | 5 | Distance and RMSD | DMscore |
| **A1R:Gi binding** | 1 (mwSuMD) | 100 ps | 3 | RMSD | SMscore |
| **GLP-1R:PF06882961** | 1 (mwSuMD) | 100 ps | 5 | Distance or RMSD (or a combination) | SMscore or DMscore |
| **GLP-1R:$G_S$ protein binding** | 3 (mwSuMD) | 200 ps | 3 | Distance or RMSD | SMscore or DMscore |
| **Gs AHD opening** | 1 (mwSuMD) | 100 ps | 3 | Distance | SMscore |
| **GLP-1R:Gs GDP unbinding** | 3 | 50 ps | 5 | Distance | SMscore |

N/A: not applicable, SuMD was not performed.

**Table S2. GLP.1R:Gs contacts during mwSuMD Gs binding simulations.**

| GLP-1R Residue | Gs Residue | Occupancy (% MD frames) |
|---|---|---|
| ASP344 (6.33) | ARG385 | 87,7 |
| ARG348 (6.37) | LEU394 | 83 |
| GLU408 (7.63) | ARG356 | 64,4 |
| LEU254 (3.57) | TYR391 | 58,1 |
| ARG419 (7.83) | ASP312 | 55,3 |
| LYS415 (7.70) | ASP312 | 47,6 |
| TYR250 (3.53) | TYR391 | 46,3 |
| LEU254 (3.57) | LEU393 | 43,7 |
| ARG419 (7.83) | HIS311 | 43,2 |
| LEU254 (3.57) | LEU388 | 42,4 |
| VAL259 (ICL2) | ARG38 | 39,4 |
| SER352 (6.41) | LEU393 | 37,4 |
| LYS351 (6.40) | LEU394 | 36 |
| PHE257 (3.60) | GLN384 | 35,2 |
| GLU262 (4.38) | ARG38 | 35 |
| ARG348 (6.37) | ARG385 | 30,8 |
| ARG348 (6.37) | LEU393 | 30,6 |
| LEU251 (3.54) | LEU393 | 29,4 |
| ILE345 (6.34) | ARG385 | 23,8 |
| LEU349 (6.38) | LEU394 | 23,2 |
| HIS171 (ICL1) | ASP312 | 22,3 |
| LYS342 (6.31) | ASP323 | 21,6 |
| PHE260 (ICL2) | ARG38 | 21,5 |
| LEU254 (3.57) | GLN384 | 20,6 |

| | | |
|---|---|---|
| ILE345 (6.34) | LEU394 | 19,9 |
| GLU262 (4.38) | TYR391 | 19,7 |
| SER261 (ICL2) | GLN35 | 19,7 |
| ARG348 (6.37) | ARG389 | 18,8 |
| LEU254 (3.57) | HIS387 | 18,6 |
| HIS171 (ICL1) | ASN313 | 17,6 |
| VAL259 (ICL2) | HIS387 | 16,8 |
| SER352 (6.41) | LEU394 | 16,7 |
| PHE257 (3.60) | HIS387 | 16,3 |
| LYS351 (6.40) | LEU393 | 16,3 |
| TYR250 (3.53) | LEU393 | 15,7 |
| LEU255 (3.58) | GLN384 | 15,6 |
| GLU412 (7.67) | ARG356 | 15 |
| LYS415 (7.70) | ASP354 | 14 |
| HIS173 (ICL1) | PHE335 | 13,9 |
| VAL259 (ICL2) | GLN35 | 13,8 |
| PHE257 (3.60) | LEU388 | 13,6 |
| HIS171 (ICL1) | ASP333 | 13,5 |
| SER258 (ICL2) | HIS387 | 13,1 |
| SER261 (ICL2) | ARG38 | 12,9 |
| ASN407 (7.62) | ARG356 | 12,5 |
| VAL259 (ICL2) | TYR391 | 12,4 |
| ARG170 (ICL1) | ASP312 | 12,3 |
| LEU255 (3.58) | LEU388 | 11,6 |
| LEU411 (7.66) | ARG356 | 11,5 |
| ARG176 (2.46) | GLU392 | 11,5 |
| GLN263 (4.39) | TRP332 | 11,5 |
| ILE345 (6.34) | LEU388 | 11,3 |
| HIS171 (ICL1) | ARG314 | 11,2 |
| VAL259 (ICL2) | ALA39 | 10,7 |
| TRP264 (4.40) | PHE335 | 10,5 |
| ARG267 (4.43) | ASP333 | 10,5 |
| HIS171 (ICL1) | TRP332 | 10,5 |
| PHE257 (3.60) | ARG38 | 10,2 |
| LEU349 (6.38) | LEU393 | 10 |
| GLN263 (4.39) | ASP333 | 10 |
| GLN263 (4.39) | ASN313 | 9,9 |
| ARG419 (7.83) | ASP354 | 9,7 |
| THR343 (6.32) | ARG385 | 9,1 |
| SER258 (ICL2) | ARG38 | 9 |
| LYS342 (6.31) | GLU322 | 8,7 |
| SER261 (ICL2) | ASP333 | 8,6 |
| LEU255 (3.58) | LEU394 | 8,6 |
| PHE260 (ICL2) | TYR391 | 8,5 |
| ASN406 (7.61) | GLU392 | 8,4 |
| SER258 (ICL2) | HIS41 | 8,4 |
| HIS171 (ICL1) | ARG337 | 8,4 |
| SER258 (ICL2) | GLN384 | 8,1 |
| ARG348 (6.37) | LEU388 | 7,9 |
| ARG267 (4.43) | ASN313 | 7,9 |
| HIS173 (ICL1) | ARG337 | 7,8 |

| | | |
|---|---|---|
| PHE260 (ICL2) | GLN35 | 7,4 |
| ARG176 (2.46) | TYR391 | 7,3 |
| ARG267 (4.43) | PHE335 | 7,2 |
| ALA256 (3.59) | GLN384 | 7 |
| VAL259 (ICL2) | HIS41 | 7 |
| CYS174 (2.44) | ARG356 | 6,5 |
| GLU408 (7.63) | GLU392 | 6,1 |
| PHE257 (3.60) | TYR391 | 6,1 |
| SER261 (ICL2) | LYS34 | 6 |
| HIS173 (ICL1) | ARG314 | 6 |
| LEU251 (3.54) | LEU388 | 6 |
| HIS173 (ICL1) | ASP312 | 5,9 |
| HIS173 (ICL1) | TRP332 | 5,9 |
| VAL259 (ICL2) | LEU55 | 5,9 |
| ASN407 (7.62) | ARG389 | 5,8 |
| HIS171 (ICL1) | GLY310 | 5,8 |
| TYR178 (2.48) | ASP312 | 5,7 |
| SER258 (ICL2) | VAL217 | 5,7 |
| SER261 (ICL2) | GLN31 | 5,7 |
| LYS342 (6.31) | THR325 | 5,6 |
| LYS342 (6.31) | ALA324 | 5,5 |
| LEU254 (3.57) | LEU394 | 5,4 |
| CYS341 (6.30) | ASP323 | 5,4 |
| TRP264 (4.40) | ARG52 | 5,3 |
| GLU408 (7.63) | ARG389 | 5,2 |
| SER258 (ICL2) | TYR391 | 5,1 |
